# Ritonavir-Induced Cellular Stress Alters Viral HBs Glycoprotein Biogenesis and Production of Infectious Hepatitis D Virions

**DOI:** 10.64898/2026.03.20.713249

**Authors:** Walid El Orch, Clémence Jacquemin, Florentin Pastor, Romain Barnault, Fanny Charriaud, Amaury Wuilbaut, Clément Nabaile, Clémentine Deleuze, Margaux Michel, Hiroshi Kadokura, Massimiliano Gaetani, Margot Begue, Eric Richard, Camille Sureau, Bernard Verrier, Christophe Ramière, Yves L. Janin, Vincent Lotteau, David Durantel, Pierre-Olivier Vidalain

## Abstract

Chronic co-infections by HBV and its satellite virus HDV are associated with a high risk of progression to cirrhosis and liver cancer, and therapeutic options for achieving a cure are still unsatisfactory. HBs is the main surface glycoprotein of both viruses, and is also massively secreted by infected hepatocytes in the form of empty subviral particles which suppress the host immune responses. This makes HBs an attractive target to develop therapeutic strategies. Here, we took advantage of the known interaction between the Large form HDV antigen (HDAg-L) and the small form of HBs (S-HBs) to develop a non-infectious, minimalistic reporter assay for the assembly and secretion of HDV particles. By screening the existing pharmacopeia for drugs that could interfere with S-HBs and HDAg-L co-secretion, we found that ritonavir and other Cytochrome P450 inhibitors affect the biogenesis of HBs and impair the production of infectious HDV virions. Mechanistically, we established that these drugs induce oxidative stress which dysregulates disulfide bond formation in the endoplasmic reticulum. As a consequence, the production of HBs, which depends on a dense network of disulfide bonds, is markedly affected as evidenced by an abnormal glycosylation profile, altered antigenic properties, and a poor expression of the largest form of HBs (L-HBs) which is essential to virus entry into target cells. This is associated with induction of the unfolded protein response, with the upregulation of CHOP/DDIT3 and key enzymes involved in the synthesis of the reducing metabolite glutathione (PHGDH, SHMT2, MTHFD2). Overall, our results indicate that alterations in redox homeostasis significantly impact HBs biogenesis, and reveal a druggable pathway that could be exploited to eliminate HDV in chronically infected patients.

**IMPACT AND IMPLICATIONS:** More effective therapies are still needed to achieve a functional cure in patients chronically co-infected by HBV and HDV. In this study, we discovered that ritonavir, along with other cytochrome P450 inhibitors, can affect the production of infectious HDV particles in human hepatocyte cultures. Mechanistically, ritonavir induces oxidative stress and the unfolded protein response in the endoplasmic reticulum, thereby altering the biogenesis of HBs, the surface glycoprotein of both viruses. This work highlights the potential benefit and mechanism of action of ritonavir and related molecules in the treatment of co-infected patients.

## INTRODUCTION

Hepatitis B, caused by Hepatitis B virus (HBV), is a major public health issue, with approximately 254 million people living with chronic infection worldwide (Fact-sheets 2024, WHO; https://www.who.int/news-room/fact-sheets/detail/hepatitis-b). It is primarily transmitted through contact with body fluids, including bloodborne, sexual, and mother-to-child transmission. HBV is an enveloped DNA virus belonging to the *Hepadnaviridae* family and *Orthohepadnavirus* genus, with a circular, partially double-stranded genome [1]. Upon infection, HBV enters hepatocytes where viral genome is converted into covalently closed circular DNA (cccDNA) in the nucleus, which serves as a template for viral mRNA transcription. Viral proteins, including the surface glycoprotein (HBs), core protein, and reverse transcriptase, are then packaged along with newly synthesized viral genomes into new virions. HBV infection can lead to either an acute or chronic trajectory. While acute infection can resolve spontaneously without long-term clinical consequences, a significant proportion of patients develop chronic infection, leading to severe complications such as cirrhosis and hepatocellular carcinoma, if left untreated. Chronically infected patients are treated with antiviral agents such as tenofovir and entecavir that block the reverse transcription of HBV genome [2,3]. Although this efficiently suppresses viral replication and prevents disease progression, no treatment currently offers a long-term cure of chronic infection.

Hepatitis D virus (HDV) is a satellite virus of HBV, requiring the presence of HBV for its propagation. Indeed, HDV relies on HBV surface glycoprotein HBs for packaging and release of new viral particles, as well as for cell entry [4]. Co-infection by these two viruses dramatically increases the risk of cirrhosis and hepatocellular carcinoma [4]. HDV infection affects approximately 5% of individuals with chronic hepatitis B, translating into nearly 12 million estimated cases worldwide [5]. HDV belongs to the *Kolmioviridae* family and *Deltavirus* genus, and has a small, circular single-stranded RNA genome, making it one of the smallest known RNA viruses infecting animals. HDV enters hepatocytes, where its RNA genome is transported to the nucleus. Genomic and antigenomic RNA are produced through a rolling-circle replication mechanism involving a ribozyme-mediated self-cleavage process. In parallel, the viral genome is transcribed by the host RNA polymerase II into mRNA, allowing the synthesis of the small and large forms of the HDV antigens, namely HDAg-S (195 amino acids) and HDAg-L (214 amino acids) [6]. Both isoforms are essential for viral replication, assembly of new viral ribonucleoproteins, and egress from the nucleus to virion assembly sites. Pegylated-interferon alpha (Peg-IFNa) is the first line treatment, but its efficacy is rather modest (around 10% sustained virologic response; SVR), and it is often poorly tolerated [7]. Bulevirtide, a viral entry inhibitor, has provided a significant therapeutic option for high-risk patients infected by HDV and has been approved in Europe [8]. The combination of Bulevertide and Peg-IFNa yields higher SVR rates [9], although this combination is yet to be recommended as best clinical practice due to a lack of a FDA approval of Bulevirtide. Further studies are needed to optimize and define the long-term benefits of these therapeutic options. In fact, novel anti-HBV/HDV drugs, ideally targeting both viruses, are needed to enrich the current therapeutic arsenal.

HBs, the envelope glycoprotein of HBV, which is also shared with HDV, consists of three surface isoforms, namely S-HBs, M-HBs, and L-HBs, all encoded by a single gene [10]. The shortest isoform, S-HBs, contains four transmembrane domains and exposes an extracellular loop harboring the immunodominant “a” determinant known as “HBsAg”. The S-HBs isoform is by far the most abundant of the three isoforms. It can self-assemble into empty subviral particles (SVPs) that outnumber infectious virions by up to 10,000-fold, and play a critical role in suppressing the immune response against HBV. The M-HBs includes the entire S region along with an N-terminal preS2 domain, which undergoes glycosylation at Asn4. The L-HBs contains an additional preS1 domain, which is myristoylated at its N-terminal glycine (Gly2), an essential modification for viral infectivity. Notably, the N-terminal portion of the preS1 domain of L-HBs directly interacts with the hepatocyte-specific sodium taurocholate co-transporting polypeptide (NTCP), thus initiating virus attachment to target cells and internalization. In HBV/HDV co-infected patients, S-HBs is mostly responsible for the assembly and secretion of HDV particles. The presence of L-HBs in HDV virions is also absolutely required for NCTP-mediated entry into hepatocytes. Therefore, disruption of these key post-translational modifications (glycosylation of S-HBs at Asn146 and/or myristoylation of L-HBs preS1 domain), can impair the infectivity of both HBV and HDV. The HDV hijacks the envelope of HBV through a specific interaction between HDAg-L and HBs. Indeed, isoprenylation of the C-terminal cysteine of HDAg-L enables its interaction with the cytosolic loops of HBs to promote the envelopment of HDV ribonucleoproteins. Specific tryptophan residues in HBs (e.g., W196, W199, W201 in S-HBs) are essential for this interaction, highlighting the intricate dependency of HDV on HBV structural components for infectivity and propagation [11]. Because of this dependence on HDAg-L farnesylation, the farnesyltransferase inhibitor lonafarnib has emerged as a promising therapeutic candidate for inhibiting HDV particle assembly and release [12]. Results of a large phase-III clinical trial have been disclosed [13]; yet approval has not been granted as off today. Combination approaches will be likely needed to achieve a high rate of HDV cure.

Additionally, nucleic acid polymers (NAPs) have been shown to interfere with HBs assembly, which inhibits the secretion of virions and SVPs [12]. Among them, REP 2139 is one of the most advanced NAPs and has been evaluated in clinical trial to treat co-infected patients. The results, yet to be confirmed by further clinical investigations, are promising although its use does suffer from a non-optimal administration route, *i.e.* intravenous infusion every week. Monoclonal antibodies targeting HBsAg are also in development, and phase-II clinical trials showed a virological response in more than 50% of co-infected patients [12]. However, larger studies are still required to confirm the efficacy of these treatments and better define the safety profile of these molecules. In any cases, it is likely that functional cure in co-infected patients will be only achieved by a multi-pronged approach combining several active molecules, in particular if a finite duration treatment is envisaged.

Therefore, to identify new drugs interfering with HBs assembly and secretion in the context of HDV infection, we have developed a minimalistic cell-based assay, which takes advantage of HDAg-L association with HBs in the endoplasmic reticulum (ER). This reporter system was then used to screen a small chemical library composed of drugs targeting metabolic pathways to identify hit molecules in the existing pharmacopeia. Characterization of these hits revealed that clinically approved Ritonavir and related compounds inhibiting CYP3A enzymes are triggering oxidative and ER stresses which lead to altered HBs folding and reduced infectivity of secreted virions. The link between CYP3A inhibition, cellular stress and HBs folding thus appeared as a potential druggable pathway, which could be exploited to interfere with the function of HBs in HBV and HDV-infected patients.

## MATERIAL AND METHODS

Experimental procedures are available in Supplementary Data

## RESULTS

### Development of an assay to identify drugs that interfere with the co-secretion of HDAg-L with S-HBs

To identify drugs that interfere with the interaction and co-secretion of S-HBs and HDAg-L, we designed a reporter cell line stably expressing both proteins, HDAg-L being tagged with the HiBiT peptide (Fig. 1A). This system enables a dual readout: the detection of HBsAg by chemiluminescent immunoassay (CLIA), and the quantification of HDAg-L *via* the HiBiT peptide. To generate this assay, the Huh7 hepatocyte cell line was first stably-transduced with a lentiviral vector encoding HDAg-L with an N-terminal HiBiT tag (HiBiT-HDAg-L). In these cells, the HiBiT signal was only detected in cell lysates but not in culture supernatants, consistent with the fact that HDAg-L alone is not secreted (Fig. 1B). When cells were transfected with a plasmid encoding S-HBs, the corresponding antigen HBsAg was significantly detected in culture supernatants by CLIA as expected (Fig. 1B, left panel). In addition, S-HBs expression resulted in the co-secretion of HiBiT-HDAg-L as assessed by the detection of the HiBiT tag in culture supernatants (Fig. 1B, middle panel). This demonstrates that when co-expressed, HiBiT-HDAg-L and S-HBs interact and are co-secreted, probably in the form of subviral particles with S-HBs at their surface and HiBiT-HDAg-L inside. In addition, we also showed that intracellular levels of HiBiT-HDAg-L were not significantly affected by the expression of S-HBs (Fig. 1B, right panel). We thus stably-transduced the Huh7 cells expressing the HiBiT-HDAg-L protein with a second lentivirus encoding S-HBs to establish a stable cell line expressing both viral proteins. For further validation of this reporter cell line, named Huh7(HiBiT-HDAg-L/S-HBs), we also tested the effect of lonafarnib which inhibits the farnesylation of HDAg-L, and thus prevents its association with S-HBs [12]. As expected, lonafarnib potently reduced HDAg-L secretion as assessed by inhibition of the HiBiT signal in culture supernatants while having no effect on S-HBs secretion or cell viability (Fig. 1C). Overall, this makes the Huh7(HiBiT-HDAg-L/S-HBs) cell line suitable for screening drugs that inhibit the shedding of S-HBs or its association of HDAg-L.

**Fig. 1.**
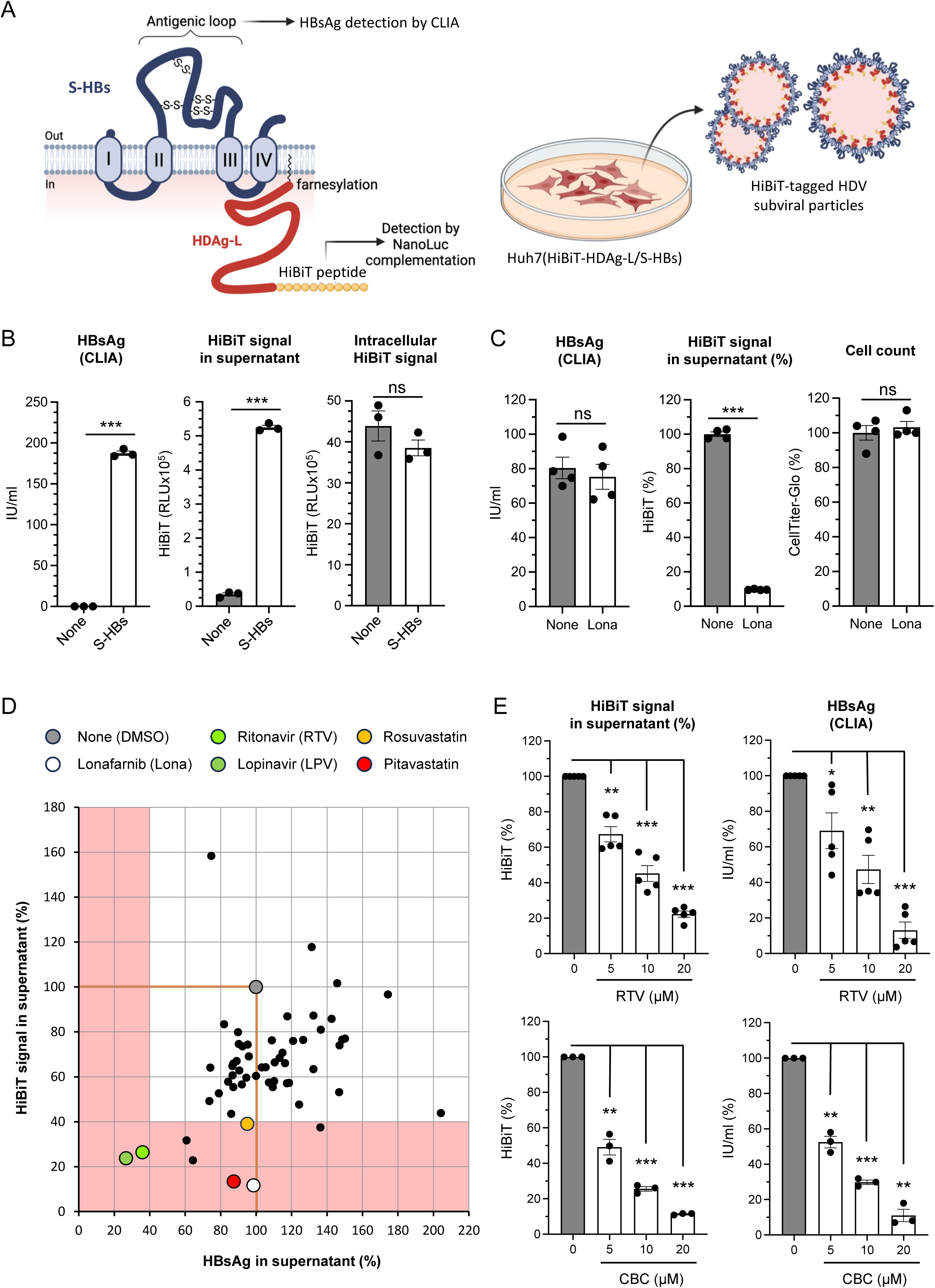
A new cell-based assay enables the identification of ritonavir and related molecules as drugs affecting the co-secretion of HDAg-L with S-HBs. **(A)** Schematic representation of the reporter cell line co-expressing the small isoform of HBV envelope glycoprotein (S-HBs) and the large delta antigen (HDAg-L) of HDV, allowing the production of HDV subviral particles (SVPs). HDAg-L is tagged with HiBiT for luminescent quantification, while S-HBs secretion is measured using a chemiluminescent immunoassay (CLIA) detecting HBsAg. **(B)** Functional validation of the Huh7-HiBiT-HDAg-L cell line showing that HDAg-L secretion in supernatant only occurs upon transfection with a plasmid encoding S-HBs. Intracellular levels of HDAg-L remain unaffected by S-HBs expression. Mean ± SEM of one experiment in triplicate; two-tailed t-test. **(C)** HDAg-L secretion is inhibited by lonafarnib, confirming that in the Huh7(HiBiT-HDAg-L/S-HBs) cell line, the farnesylation of HDAg-L is essential for its association and co-secretion with S-HBs. Mean ± SEM of one experiment in quadruplicate; two-tailed t-test. **(D)** Screening of 80 molecules from a metabolic compound library using the Huh7(HiBiT-HDAg-L/S-HBs) cell line. Data are expressed as percentages relative to DMSO for both HDAg-L (HiBiT signal) and HBsAg (CLIA) detected in supernatants. Red lines correspond to the signal in DMSO control wells. **(E)** Dose–response validation of the effects of ritonavir (RTV) and cobicistat (CBC) using the same reporter cell line. Mean ± SEM of independent experiments (raw values were first normalized to DMSO control in each experiment). One sample t-test using 100% as reference and Holm-Šidák correction for multiple testing.

### Ritonavir and related compounds interfere with the co-secretion of HDAg-L and S-HBs

To identify cellular pathways interfering with the co-secretion of S-HBs and HDAg-L, we used the Huh7(HiBiT-HDAg-L/S-HBs) cell line to screen a library of 493 metabolism-related compounds. Cells were treated with the compounds at 10 μM and supernatants were harvested 48 h later to quantify both HiBiT-HDAg-L and HBsAg. The toxicity of tested compounds was assessed in parallel for counter screening, and those showing >25% toxicity as compared to control were discarded. Compounds inhibiting the secretion of HDAg-L and/or HBsAg by >60% were selected for further evaluation. Finally, lonafarnib was used as a positive control for inhibition of HDAg-L secretion, while DMSO was used as a negative control in each screening plate. Fig. 1D shows the activity of 80 compounds from one of the screening plates where several compounds retained our attention. First, two of the molecules that reduced the secretion of HDAg-L without affecting HBsAg were statins (pitavastatin and rosuvastatin). Indeed, inhibition of HMGCR (3-hydroxy-3-methyl-glutaryl-coenzyme A reductase) by these drugs inhibits the mevalonate pathway and therefore impairs the synthesis of farnesyl-PP which is required for the farnesylation of HDAg-L [14]. Besides, two of the best hits were structurally-related peptidomimetics and HIV protease inhibitors: ritonavir and lopinavir. These molecules are also well known for interacting with cytochrome P450 enzymes involved in the oxidation of xenobiotics, especially CYP3A4 and other members of the CYP3A subfamily [15]. For this reason, ritonavir is often co-administered with other drugs to inhibit their degradation in the liver, thus increasing their plasma concentration and half-life in patients. Surprisingly, ritonavir and lopinavir inhibited both HDAg-L and HBsAg parameters (Fig. 1D), suggesting they interfere with S-HBs overall biogenesis (i.e.: assembly or/and secretion), warranting further mechanistic investigations.

Following the initial screening, we first validated the activity of these two molecules using the same reporter cell line, with dose-response concentrations ranging from 5 to 20 µM. Results confirmed that ritonavir (RTV) and lopinavir (LPV) inhibit levels of HDAg-L and HBsAg in culture supernatants, as assessed by HiBiT quantification and CLIA, respectively (Fig. 1E, top panels; and Fig. S1A). In addition, ritonavir and lopinavir did not show significant toxicity in cell cultures (Fig. S1B). Several structural analogues of ritonavir with a more favorable safety profile have been developed as CYP3A4 inhibitors and pharmacokinetic enhancer. This includes cobicistat, which does not inhibit HIV protease, but is widely used in combinations with HIV protease inhibitors to improve their half-life [16]. We thus tested cobicistat (CBC), and a dose-response inhibition of HDAg-L and HBsAg parameters was observed in treated cells (Fig. 1E, bottom panels) without significant cellular toxicity of this molecule (Fig. S1B). Overall, this shows that ritonavir and two structurally-related drugs have similar effects on HDAg-L and S-HBs co-secretion.

### Ritonavir inhibits the production of infectious HDV particles in primary human hepatocytes

Based on these results, we then studied the effects of ritonavir on HDV replication. To explore this, we analyzed the effect of ritonavir on freshly isolated human hepatocytes (PHH) co-infected with HBV and HDV. Cells were cultured for six days to fully establish infection before starting the treatment (Fig. 2A)[17]. Cells were treated for 6 days with cell culture medium containing ritonavir (10 μM) or not, with medium replacement after three days of culture. Finally, cells and culture supernatants were harvested to quantify viral growth. Ritonavir was not toxic as assessed by the absence of Lactate Dehydrogenase (LDH) release in culture supernatants (Fig. S2A), and showed no effect on HBV or HDV replication as assessed by RT-qPCR on cell lysates (Fig. 2B). We also evaluated the effect of ritonavir on HBV and HDV genomes in PHH supernatants as well as the secretion of HBsAg. As shown in Figure 2C, the secretion of HBV and HDV genomes was not affected by ritonavir. In contrast, when quantifying HBsAg by CLIA, a decreased signal was observed in the supernatants of PHH treated with ritonavir (Fig. S2B). This suggests that ritonavir alters the antigenic properties of S-HBs, preventing detection of the antigenic loop by CLIA. Indeed, the antigenicity and functionality of HBsAg are highly dependent on its folding, and in particular on the disulfide bonds between cysteine residues that stabilize its conformation and participate in virus entry [18,19,10]. As this could reduce the infectivity of secreted virions, we evaluated the presence of infectious HDV particles in the supernatants of PHH by re-infecting differentiated HepaRG cells (dHepaRG), a well-characterized *in vitro* model that supports HDV infection [20,21]. Supernatants from PHH, diluted at 1/60^th^, were thus applied to fresh dHepaRG cells. After 6 days of culture, HDV genome in dHepaRG cells was quantified by RT-qPCR (Fig. 2D). Results showed that ritonavir strongly (>90%) reduced the amount of infectious HDV particles detected in the supernatants of PHH. It is important to note that we verified that the presence of ritonavir in the diluted PHH supernatants cannot directly block the entry of HDV into dHepaRG cells (Fig. S3)[22]. Therefore, ritonavir does not affect viral replication *per se* or the shedding/release of HDV genome, but affects S-HBs folding and decreases HDV virion infectivity.

**Fig. 2.**
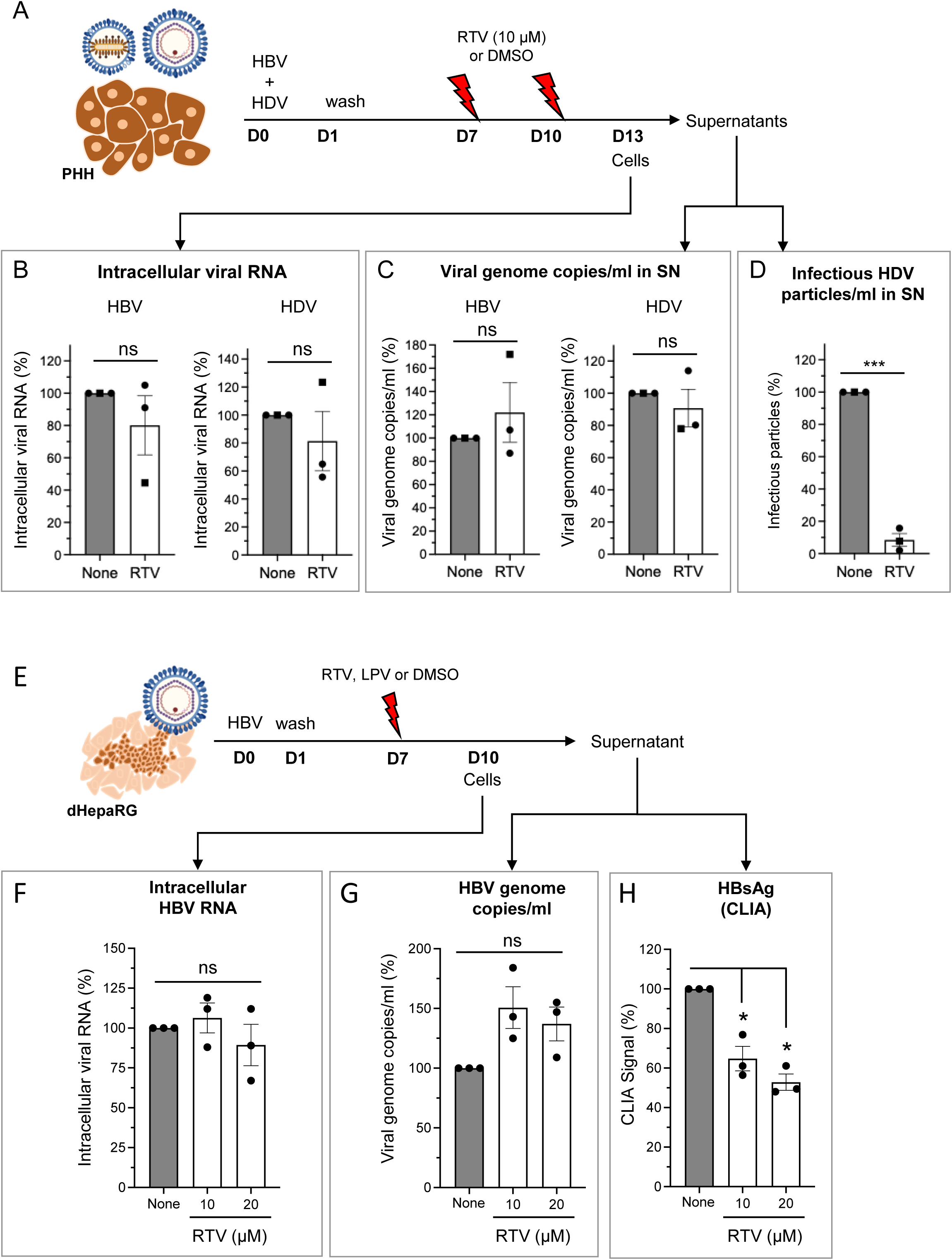
Ritonavir disrupts the production of infectious HDV particles in PHH and alters HBs biogenesis in HBV-infected dHepaRG cells. **(A)** Schematic representation of the experimental workflow used for testing the effect of ritonavir (10 μM) on PHH co-infected by HBV and HDV. **(B)** Intracellular HBV and HDV RNAs quantified by RT-qPCR, normalized to the PrP housekeeping gene. **(C)** Quantification of HBV and HDV genome copies in culture supernatants quantified by qPCR and RT-qPCR, respectively. **(D)** The presence of infectious viral particles in culture supernatants was determined by infection of fresh dHepaRG cells. PHH were directly isolated from human liver resections (black circles; two donors) or amplified in the liver of TK-NOG mice before purification (black square; HepaSH®; [56]). **(E)** Experimental workflow used to investigate the impact of ritonavir in HBV-infected dHepaRG cells. Differentiated HepaRG cells were infected with HBV and treated with ritonavir for 3 days. **(F-H)** Effect of ritonavir on HBV infection. **(F)** Intracellular HBV RNAs quantified by RT-qPCR, normalized to human RPL13A expression used as housekeeping gene. **(G)** HBV genome copies in culture supernatant quantified by qPCR using a standard curve. **(H)** HBsAg level quantified by CLIA in culture supernatant. Mean ± SEM of independent experiments (raw values were first normalized to DMSO control in each experiment). One sample t-test using 100% as reference and Holm-Šidák correction for multiple testing.

### Ritonavir alters HBs biogenesis in HBV-infected dHepaRG cells

To confirm the effect of ritonavir on HBsAg independently of HDV infection, we analyzed its effect on cells infected by HBV alone. We used differentiated HepaRG cells (dHepaRG), a well-characterized *in vitro* model that supports HBV infection [21]. These cells were infected with HBV, and cultured for seven days to allow the infection to be fully established (Fig. 2E). Treatment was added and after three days, culture supernatants and cells were harvested. Quantification of lactate dehydrogenase (LDH) activity in culture supernatants showed that ritonavir was not toxic (Fig. S4). Ritonavir had no effect on HBV replication as assessed by RT-qPCR on cell lysates (Fig. 2F). Secretion of HBV genome (*i.e.*: viremia), determined by qPCR on culture supernatants, was not significantly affected (Fig. 2G). In contrast, HBsAg detection by CLIA was reduced by ritonavir (Fig. 2H). Similar results were obtained with lopinavir (Fig. S5). This divergence between the secretion of DNA-containing HBV particles and the reduced CLIA signal is consistent with an impaired folding of HBs.

### Ritonavir alters the conformation and glycosylation profile of S-HBs

To further demonstrate that ritonavir and related molecules primarily alter the folding of S-HBs, we established a novel Huh7 cell line that stably expresses S-HBs directly fused to the HiBiT peptide at its C-terminus (Fig. 3A). This cell line, named Huh7(S-HBs-HiBiT), enables precise quantification of secreted S-HBs through detection of the HiBiT tag in culture supernatants. In addition, HBsAg, whose detection by antibodies depends on the folded antigenic loop, can be quantified in supernatants by CLIA. We thus treated Huh7(S-HBs-HiBiT) cells with increasing concentrations of ritonavir, cobicistat or lopinavir for 48 h to assess their impact on S-HBs. The effect of these drugs on the HiBiT signal was rather limited, and did not reach statistical significance with ritonavir, even at the highest concentration (Fig. 3B, left panel). Inhibition reached a maximum of ∼50% with cobicistat and lopinavir (Fig. 3C left panel and Fig. S6A). In the meantime, the detection of HBsAg by CLIA was very significantly reduced in the presence of these drugs. Inhibition was >80% with ritonavir at 20 µM, whether the results were expressed as a percentage of the raw CLIA signal (Fig. 3B, middle panel), or as a percentage HBsAg concentration in IU/ml after transposition using the kit’s dose-response curve (Fig. S6B). Even greater inhibitory effects were obtained with cobicistat and lopinavir (Fig. 3C middle panel and Fig. S6A, respectively). Again, cell viability was not significantly affected by these three drugs (Fig. 3B-C right panels and Fig. S6A-B). Overall, results demonstrate that these drugs alter the conformation of S-HBs, preventing the recognition of HBsAg by antibodies specific for the antigenic loop used in CLIA, while the effect on S-HBs secretion ranges from low (ritonavir) to moderate (lopinavir and cobicistat).

**Fig. 3.**
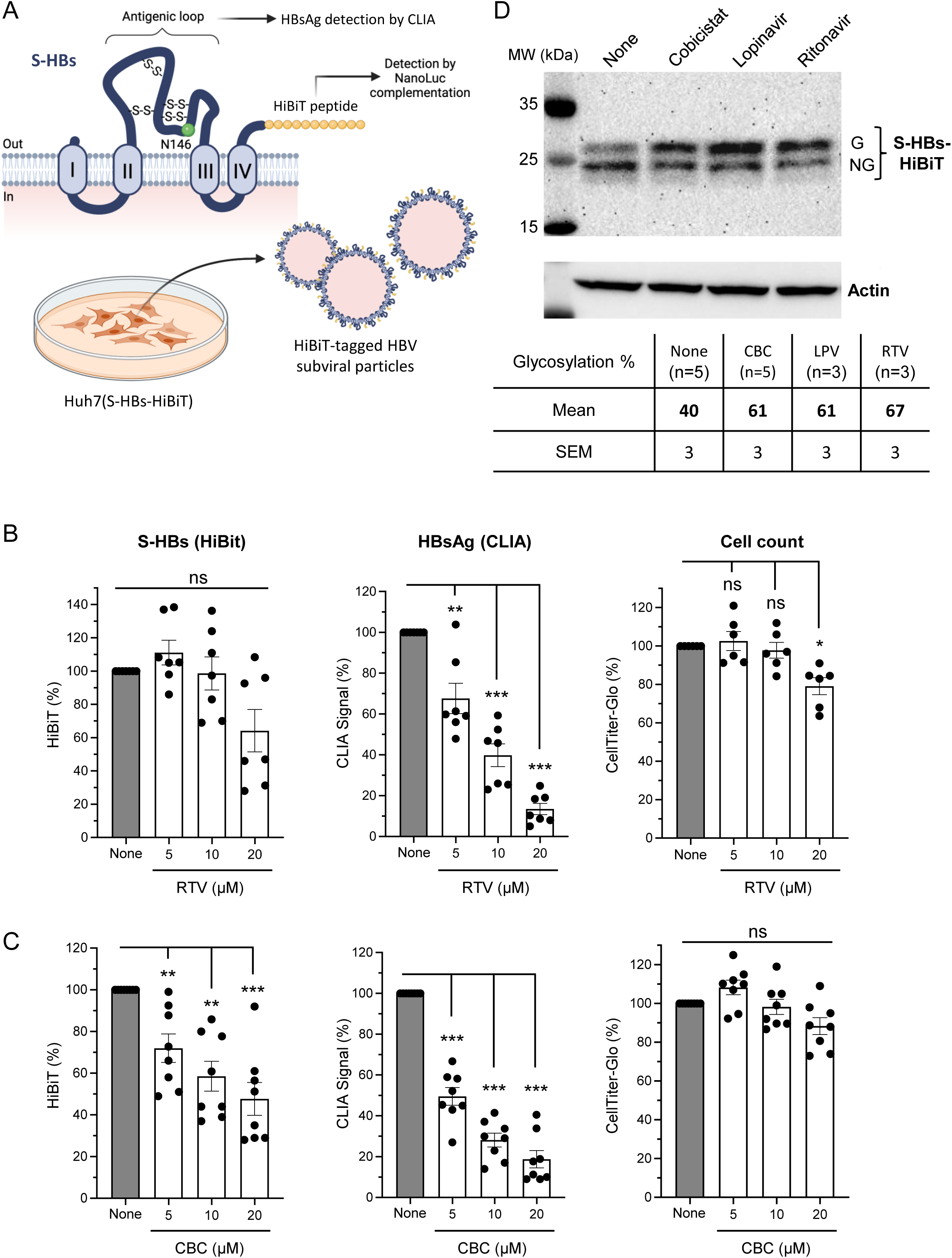
Effect of ritonavir and related compounds on S-HBs antigenicity and glycosylation. **(A)** Schematic representation of the Huh7 reporter cell line stably expressing HBV small surface antigen (S-HBs) C-terminally fused to a HiBiT tag. This system enables dual readouts: quantification of S-HBs secretion by HiBiT quantification using luminescence, and assessment of S-HBs antigenicity by CLIA assay (HBsAg quantification). **(B)** After 48 h of culture, the effect of ritonavir treatment on S-HBs secretion (HiBiT signal), HBsAg level (CLIA signal), and cell viability (CellTiter-Glo signal) were determined. **(C)** Effect of cobicistat on the same parameters, shown in the same order and format as in panel B. Data in (B) and (C) correspond to the mean ± SEM of independent experiments (raw values were first normalized to DMSO control in each experiment). One sample t-test using 100% as reference and Holm-Šidák correction for multiple testing. **(D)** Effect of ritonavir and related compounds cobicistat and lopinavir at 20 μM on S-HBs glycosylation. The percentage of the glycosylated form for each condition is shown below in the table (Mean ± SEM). It corresponds to the intensity of the signal in the upper band divided by the sum of the intensities of the upper and lower bands. Actin level is used as loading control.

To confirm the effect of these drugs on the folding of S-HBs, we characterized their impact on its glycosylation profile. Indeed, S-HBs is glycosylated at position N146 in the antigenic loop. The folding of S-HBs makes this glycosylation site only partially accessible, leading to ratios of 1/2 between glycosylated and non-glycosylated monomers of S-HBs. When mutations are introduced at positions C147 and C149, key disulfide bonds involved in S-HBs folding cannot be formed and the corresponding mutant is more efficiently glycosylated [18,19]. We thus analyzed by western-blot the fraction of glycosylated and non-glycosylated monomers of S-HBs expressed in the Huh7(S-HBs-HiBiT) cell line. As shown in Figure 3D, the glycosylation rate of S-HBs in DMSO-treated control cultures is close to 40% as expected. However, the fraction of glycosylated S-HBs increased to ∼61-67% when cells were treated with ritonavir, lopinavir or cobicistat (Fig. 3D). These results suggest that the glycosylation site at position N146 is made more accessible in cells treated with these drugs, supporting an effect of ritonavir-related compounds on the folding of S-HBs.

We then assessed the effect of ritonavir and lopinavir on HBs in a cellular model expressing the small (S), the medium (M) and the large (L) forms of HBs. To this end, Huh7 cells were transfected with a plasmid expressing all three forms of HBs under the control of the endogenous HBV promoter (pT7HB2.7). Cells were then treated for 48 h, and HBsAg secreted in culture supernatant was first quantified by CLIA (Fig. 4A). Both ritonavir and lopinavir strongly reduced the CLIA signal, in a dose dependent manner, supporting a major defect in the secretion of correctly folded HBs. We then analyzed by western blot the expression of HBs and its glycosylation profile (Fig. 4B). Ritonavir and lopinavir at 10 µM led to a significant increase in the glycosylated form of S-HBs, accompanied by a marked reduction in the non-glycosylated form. At 20 µM, S-HBs was mostly expressed under its glycosylated form, and overall expression of S-HBs was clearly reduced when compared to control. Interestingly, the M and L forms of HBs were also strongly suppressed at 10 µM and were undetectable at 20 µM. Although M-HBs is not critical to virion assembly and secretion, the net loss and/or relative loss of incorporation of L-HBs will have profound implications for viral infectivity as it plays a critical role in receptor binding and virion entry (10). The disappearance of this M and L-HBs isoforms suggests that ritonavir-related compounds alter the folding and glycosylation pattern of HBs, but may also modify the stoichiometry of the viral envelope on HDV particles, thus reducing their infectivity.

**Fig. 4.**
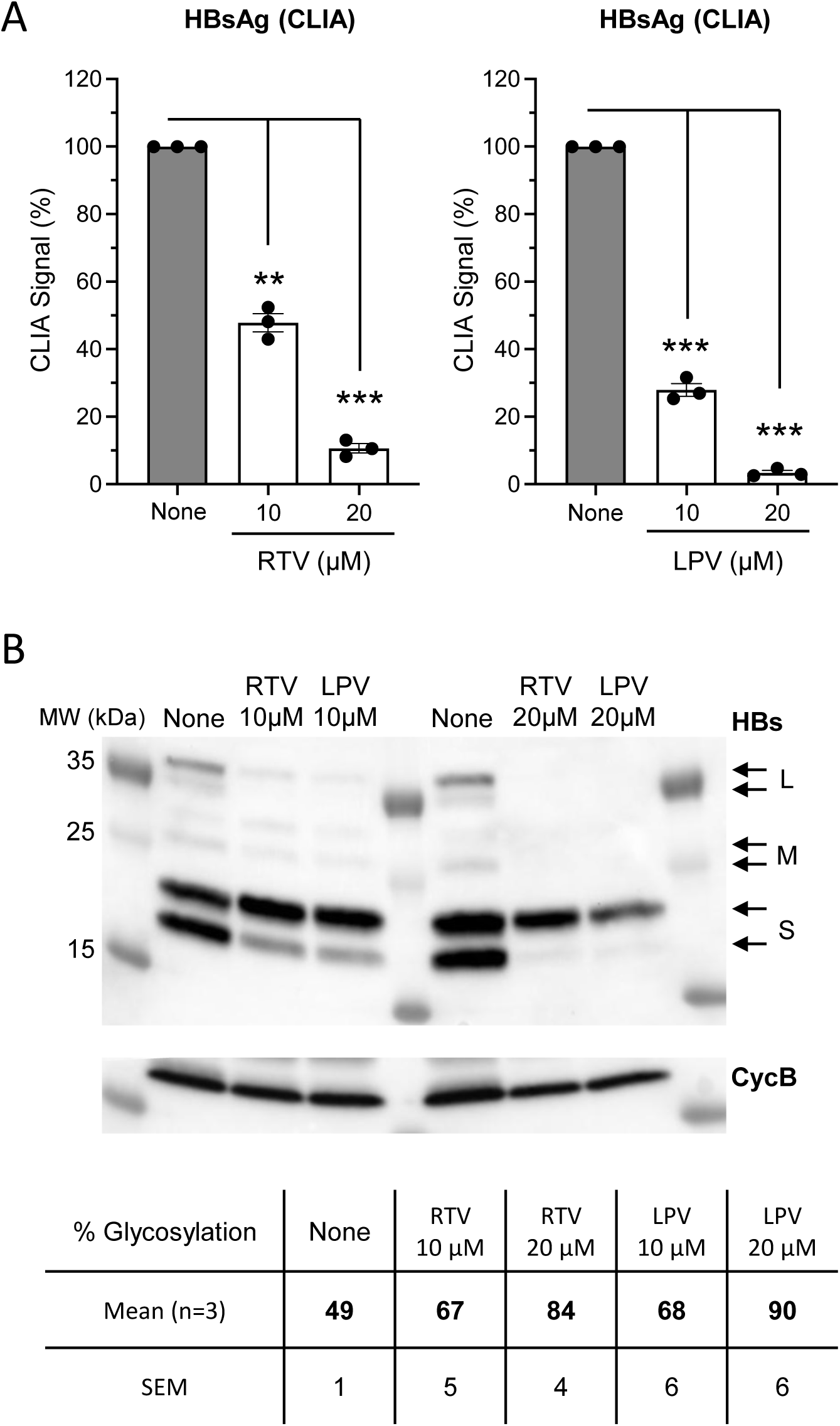
Effect of ritonavir and lopinavir on the antigenicity, glycosylation and expression of HBs isoforms expressed by the pT7HB2.7 plasmid transfected in Huh7 cells. **(A)** Dose-dependent effect of ritonavir and lopinavir on HBsAg level in the supernatant of Huh7 cells transfected with pT7HB2.7. Mean ± SEM of independent experiments (raw values were first normalized to DMSO control in each experiment). One sample t-test using 100% as reference and Holm-Šidák correction for multiple testing. **(B)** Impact of ritonavir and lopinavir on HBs expression and glycosylation for all three isoforms (L, M and S). Cyclophilin B was used as a loading control. The percentage of glycosylated S-HBs for each culture condition is shown in the table below (Mean ± SEM of three independent experiments).

### The CYP3A4 inhibitor ketoconazole also alters the conformation and glycosylation of S-HBs

We wanted to further investigate the molecular mechanisms underlying the effect of ritonavir and related compounds on S-HBs. Because of its ability to block CYP3A4 and other related CYP3A enzymes involved in xenobiotic metabolism, ritonavir is extensively used for improving drug pharmacokinetics [15]. To determine whether CYP3A inhibition accounts for the effect of ritonavir and related molecules (lopinavir and cobicistat) on S-HBs, we took advantage of ketoconazole, a well-characterized CYP3A inhibitor whose chemical structure is totally different from ritonavir. To determine its effect on S-HBs, we treated Huh7(S-HBs-HiBiT) cells with ketoconazole for 48 h and then analyzed HBsAg secretion. Ketoconazole efficiently reduced the CLIA signal while the effect on the HiBiT signal was quite limited and did not reach statistical significance, strongly suggesting that S-HBs folding is altered (Fig. S7A, left and middle panels). In addition, cell viability was not significantly affected by this drug (Fig. S7A, right panel). We also analyzed cellular lysates by western-blot to determine the fraction of glycosylated and non-glycosylated monomers of S-HBs. As shown in Figure S7B, the fraction of glycosylated S-HBs in proportion to the concentration of ketoconazole, was consistent with results obtained with ritonavir and related molecules. As two structurally unrelated CYP3A inhibitors have the same phenotype, these results strongly suggest that CYP3A inhibition is responsible for the observed effect of these drugs on S-HBs.

### Transcriptomic data reveal the induction of an endoplasmic reticulum stress in ritonavir-treated cells

We sought to further investigate the molecular mechanisms underlying the effect of ritonavir and related compounds on HBs. To decipher the cellular response to these drugs, we conducted a transcriptomic analysis of Huh7 cells following treatment with ritonavir or lopinavir. To this end, Huh7 cells were treated for 48 h, and RNA transcripts were analyzed by next generation sequencing. Differentially expressed genes (DEGs) were identified using the following criteria: |log_₂_FC| > 0.5 or 1, and adjusted p-value < 0.05 (Table S1). As shown in Figure 5A, 344 genes had their expression upregulated by ritonavir, lopinavir, or both. The subset of 131 genes whose expression was induced by both ritonavir and lopinavir was analyzed for functional annotation enrichment with the DAVID analysis platform using Uniprot keywords, Gene Ontology terms associated with Biological Processes (BP) and KEGG pathway annotation (Fig. 5B). Ritonavir and lopinavir were found to induce genes associated with endoplasmic reticulum (ER) stress, including transcription factors (DDIT3/CHOP, CEBPB and ATF3), regulators of the Unfolded Protein Response (UPR) pathway (DNAJB9, DNAJC10, FICD), and effectors controlling apoptosis (NIBAN1, BBC3/PUMA). This observation is consistent with previous reports showing the induction of ER stress by HIV protease inhibitors in different cellular models [23–27].

**Fig. 5.**
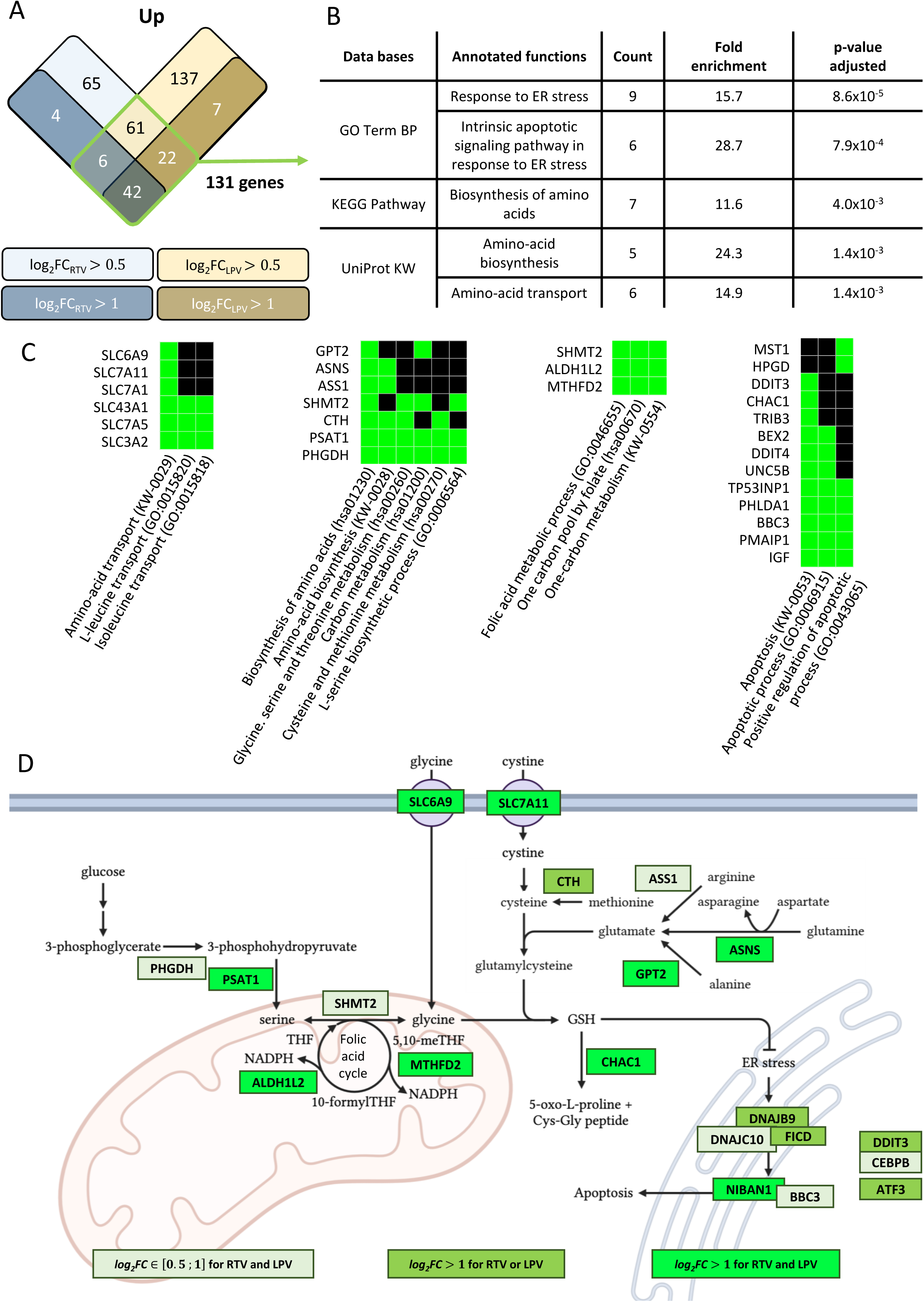
Transcriptomic analysis of upregulated genes in Huh7 cells treated with ritonavir or lopinavir. **(A)** Venn diagram showing the overlap of upregulated genes in Huh7 cells treated for 48 h with ritonavir or lopinavir. Differential expression was defined by a threshold of log_₂_FC > 0.5 or 1 and an adjusted *p*-value < 0.05. **(B)** Functional enrichment analysis of upregulated genes using the DAVID platform, based on UniProt keywords (KW), Gene Ontology (GO) Biological Process (BP) terms, and KEGG pathway annotations. **(C)** Clustering of enriched functional annotations from the three databases (UniProt, GO-BP, KEGG), highlighting the major functional categories and the genes associated with each cluster. **(D)** Schematic showing a subset of upregulated genes selected from the clusters presented in (C) and their involvement in the serine/glycine pathway, folic acid cycle, glutathione synthesis and ER stress leading to apoptosis.

Genes involved in amino acid biosynthesis and transport were also induced in ritonavir and lopinavir-treated cells (Fig. 5B). This analysis was further refined by clustering the enriched functional annotations from the three databases with DAVID, while highlighting the genes involved in these different functions (Fig. 5C). The first two clusters were made up of genes involved in amino acid transport and synthesis, respectively. As shown in Figure 5D, several genes in these two clusters contribute to the biosynthesis of glycine and cysteine, two amino acids required for the production of glutathione (GSH), a reducing metabolite essential for the elimination of xenobiotics (Phase-II metabolism) and for the maintenance of the cell’s redox status, which is particularly important in the control of protein folding in the ER. Indeed, the import of GSH to the ER is thought to buffer the activity of protein disulfide isomerases (PDIs) in this compartment and to prevent the formation of erroneous disulfide bonds in ER client proteins [28,29]. Transporters of glycine and cystine, the oxidized derivative of cysteine, were also upregulated by ritonavir and lopinavir. A third cluster of genes is composed of SHMT2, MTHFD2 and ALDH1L2. SHMT2 catalyzes the conversion of serine to glycine, but is also involved with MTHFD2 and ALDH1L2 in the folic acid cycle which, among other functions, produces NADPH to prevent oxidative stress and maintain the pool of GSH via the activity of glutathione reductase [30]. Finally, it should be highlighted that PHGDH, PSAT1, SHMT2, MTHFD2 and ALDH1L2 have all previously been associated with the cellular response to Tunicamycin- and Thapsigargin-induced ER stress, and have been assigned to the UPR regulon [31]. Overall, this transcriptomic signature shows strong markers of ER stress and induction of the UPR pathway in cells treated with ritonavir and lopinavir. A fourth group of genes was related to apoptosis regulation, consistent with induction of cellular stress.

Interestingly, we also identified 65 genes whose expression was significantly downregulated by both ritonavir and lopinavir (Fig. 6A), with functional annotations associated with glycolysis (Fig. 6B-C). Indeed, and as shown in Figure 6D, several enzymes involved in glycolysis had their expression repressed, particularly in downstream steps of this pathway. As glycolysis is linked to the pentose phosphate pathway and the folic acid cycle (Fig. 6D), both of which contribute to maintaining the NADPH pool, modulation of glycolytic enzymes could favor these two pathways that are essential for controlling the cell’s redox status and for the elimination of xenobiotics (Phase-I and II metabolism).

**Fig. 6.**
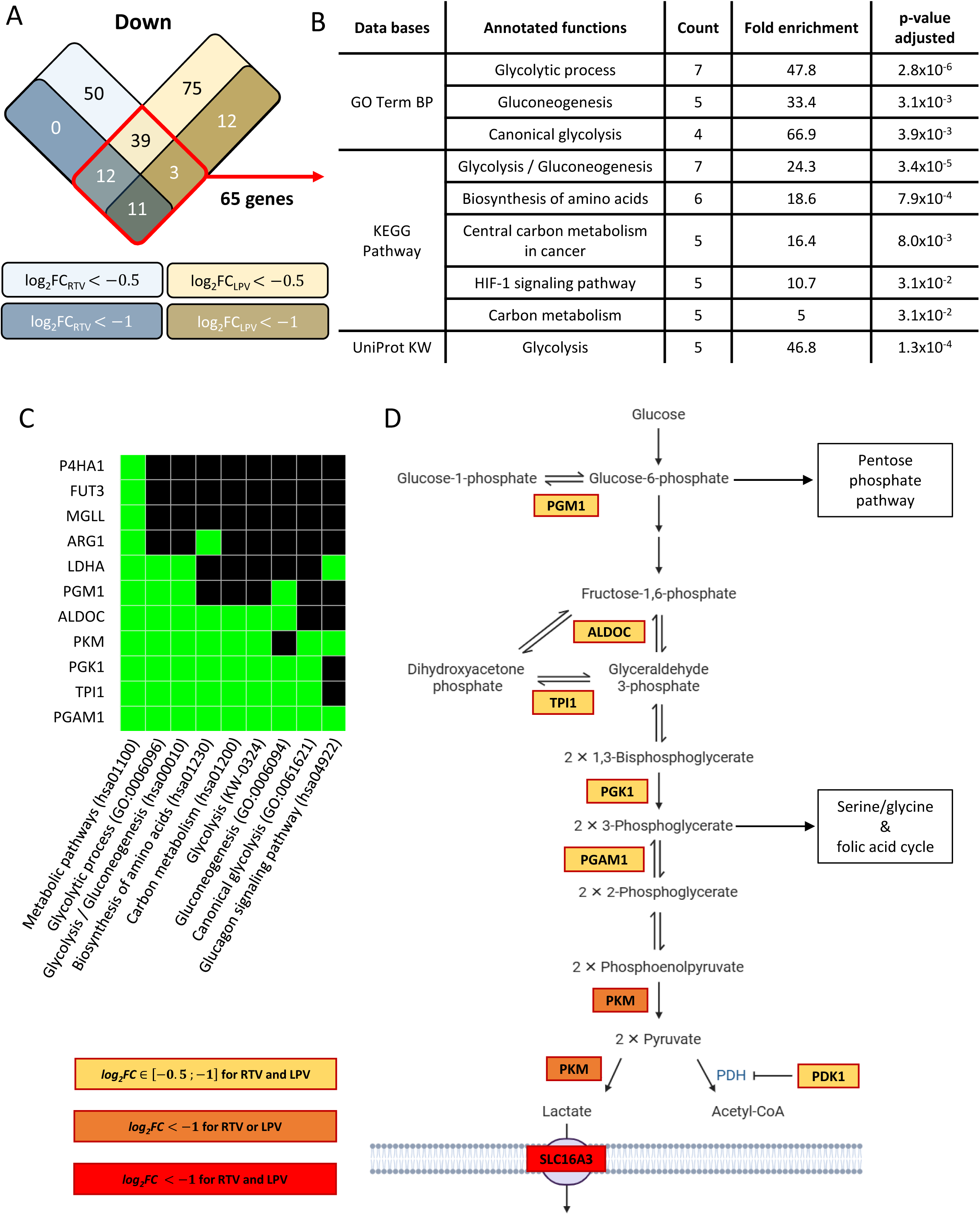
Transcriptomic analysis of downregulated genes in Huh7 cells treated with ritonavir or lopinavir. **(A)** Venn diagram showing the overlap of downregulated genes in Huh7 cells treated for 48 h with ritonavir or lopinavir. Differential expression was defined by a threshold of log_₂_FC < −0.5 or −1 and an adjusted *p*-value < 0.05. **(B)** Functional enrichment analysis of downregulated genes using the DAVID platform, based on UniProt keywords (KW), Gene Ontology (GO) Biological Process (BP) terms, and KEGG pathway annotations. **(C)** Clustering of enriched functional annotations from the three databases (UniProt, GO-BP, KEGG), highlighting the major functional categories and the associated genes. **(D)** Schematic showing a subset of downregulated genes selected from the cluster presented in (C) and their involvement in glycolysis.

Finally, to support transcriptomic data showing the induction of ER stress by ritonavir, we confirmed by RT-qPCR the upregulation of CHOP/DDIT3, which is a master activator of the UPR regulon in response to ER stress (Fig. S8A). The induction of PHGDH, SHMT2 and MTHFD2, three factors recently assigned to the UPR regulon [30–33], was also validated with this technique (Fig. S8B). These results confirm the transcriptomic data establishing the induction of the ER stress response in treated cells. This observation is consistent with a misfolding of S-HBs which could lead to a functional defect of this viral glycoprotein.

### Ritonavir induces oxidative stress as assessed by ophthalmic acid level and GSH/GSSG ratio

As many of the genes induced by ritonavir and lopinavir are metabolic enzymes, we also used untargeted metabolomic analysis performed by LC-MS/MS to document the effect of ritonavir on cellular metabolism. Of a total of 151 metabolites identified with a high level of confidence (annotation level 1 and 2a), 17 metabolites had their expression reduced in the presence of ritonavir (Fig. 7A, left panel). Decreased expression of carnitine, acylcarnitines and glycerophosphocholine suggests that lipid metabolism is dysregulated in ritonavir-treated cells. This could be linked to a change in mitochondrial activity, as suggested by decreased levels of two metabolites of the citric acid cycle, citric acid and cis-aconitic acid, as well as glycerol 3-phosphate and NADH, which supply the respiratory chain with electrons via the glycerol 3-phosphate shuttle. In addition, the expression of several glycolysis metabolites was decreased, consistent with transcriptomic data showing a modulation of multiple enzymes in this pathway (Fig. 6D). Overall, the results demonstrate profound metabolic reprogramming in cells treated with this drug. Most interestingly, GSH expression decreased in ritonavir-treated cells (Fig. 7A, left panel), consistent with an increased consumption of this metabolite to manage xenobiotic metabolism, oxidative stress, ER stress or all three combined.

**Fig. 7.**
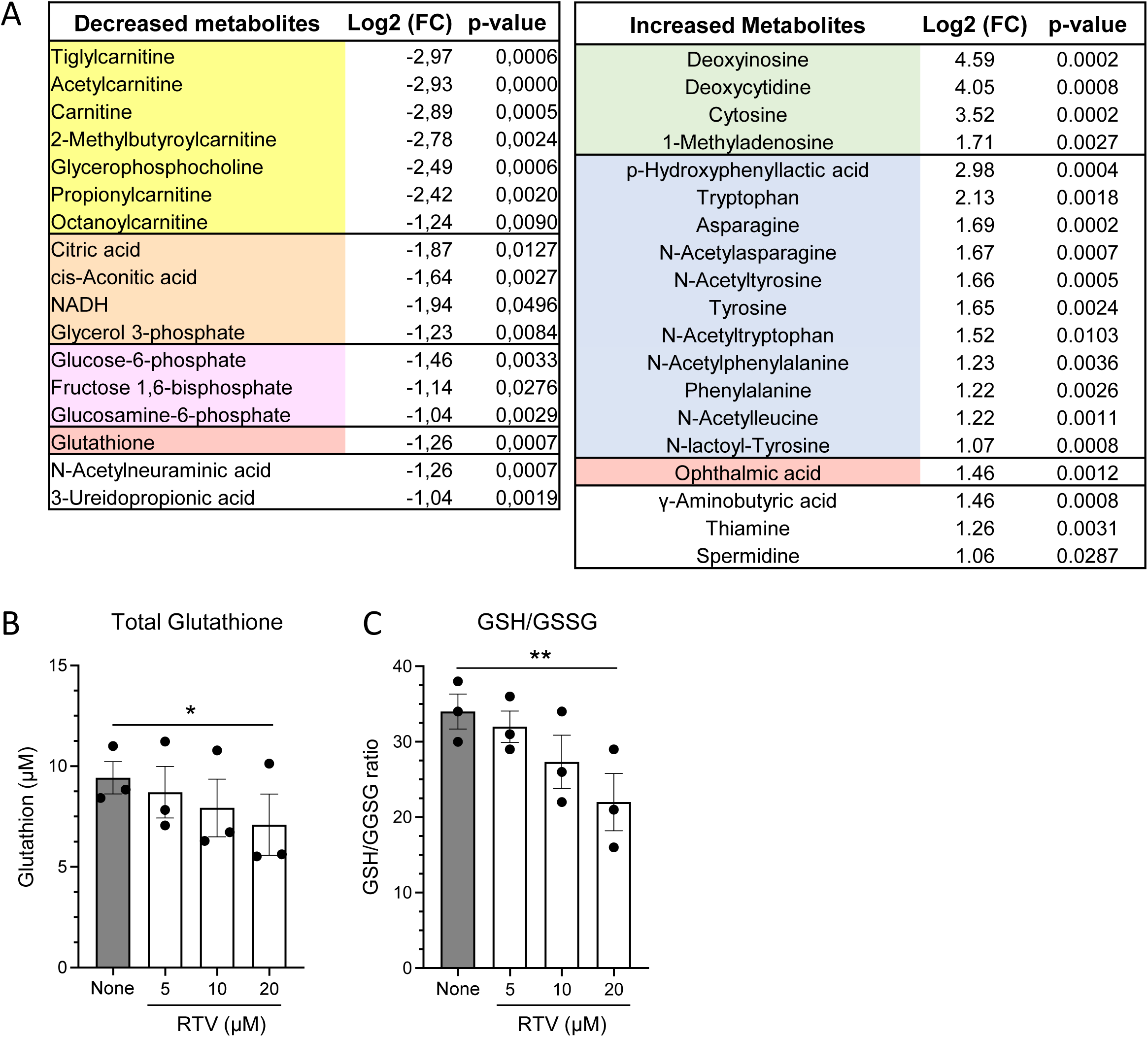
Metabolomic profiling of Huh7 cells treated with ritonavir. **(A)** Table showing metabolites whose expression was found to be increased (left) or decreased (right) in Huh7 cells treated for 48h with ritonavir. Metabolomic analysis performed by untargeted LC-MS/MS (presented metabolites are limited to annotation levels 1 and 2a; |log2FC| >1; p-value < 0.05). **(B–C)** Total glutathione (GSH+GSSG) and GSH/GSSG ratio quantified by enzymatic assay in Huh7 treated with DMSO or ritonavir for 48 h (Mean ± SEM of three independent experiments; one-way ANOVA).

We also identified 19 metabolites whose expression was increased (Fig. 7A, right panel). Several nucleotides and their metabolites showed high expression levels in ritonavir-treated cells, which could reflect an activation of the DNA damage and repair machinery and, more generally, nucleic acid degradation. The expression of several amino acid, especially aromatic ones and their N-acetylated derivatives, was also increased. Approximately 80% of proteins in humans are acetylated at their N-terminus, making it the most common protein modification [34]. As N-acetylated amino acids released during protein degradation must be processed by aminoacylases before being reincorporated into nascent proteins, the accumulation of N-acetylated amino acids could reflect increased cellular protein turnover [35]. Most interestingly, we also identified ophthalmic acid (OPH), a metabolite produced by the same pathway and similar to GSH but without the thiol group. Although the exact function of OPH remains elusive, its increased synthesis has been well documented during oxidative stress as a consequence of activation of the GSH biosynthesis pathway [36,37]. This suggests that ritonavir induces a cellular stress response leading to activation of the GSH biosynthesis pathway, in agreement with transcriptomic data (Fig. 5D).

The increased OPH levels and decreased GSH levels detected by LC-MS/MS suggest that ritonavir induces oxidative stress in Huh7 cells, which could explain the ER stress and the activation of the UPR pathway [24–26]. To explore this possibility, we first confirmed the reduced expression of total glutathione in ritonavir-treated cells using an enzymatic assay measuring both GSH and its oxidized form GSSG (Fig. 7B). Finally, another enzymatic assay was used to selectively quantify GSH and GSSG in order to determine the GSH/GSSG ratio which reflects the cell’s redox status. As shown in Figure 7C, this ratio decreased in proportion to the concentration of ritonavir added in culture medium, demonstrating the induction of oxidative stress.

### Oxidative stress is responsible for the effect of ritonavir on S-HBs

As indicated above, oxidative stress could have a direct effect on the formation of disulfide bonds in the ER. To quantify this effect, we used a modified firefly luciferase containing eight additional cysteine residues (FLuc∗)[38]. When FLuc∗ is targeted to the ER by a signal peptide (Calr (40 aa)-FLuc∗), disulfide bonds can form due to the oxidative environment (Fig. 8A, left panel). This leads to luciferase misfolding and low enzymatic activity. Calr (40 aa)-FLuc∗ can therefore be used as a molecular probe to detect redox changes in the ER, and determine whether ritonavir induces an oxidative stress promoting the formation of disulfide bonds. As the enzymatic activity of Calr (40 aa)-FLuc∗ is relatively modest compared to wild-type firefly luciferase, the plasmid encoding this reporter protein was transfected in HEK-293T cells instead of Huh7 cells to achieve protein expression levels sufficient to quantify the luminescent signal. As a control, we used Met-FLuc∗ that is expressed in the cytosol where the reducing environment favors its proper folding, making luciferase activity insensitive to variations of redox status in the ER [38]. After transfection of the plasmids expressing these reporter proteins, cells were treated with ritonavir, β-mercaptoethanol (β-ME) as reducing agent or both combined. When FLuc∗ was targeted to the ER (Calr (40 aa)-FLuc∗), its activity increased in the presence of β-ME, in agreement with a better folding of the enzyme in reducing conditions (Fig. 8A, middle panel). Furthermore, FLuc∗ activity decreased in the presence of ritonavir alone, and this effect was reverted by the addition of β-ME. In contrast, ritonavir or β-ME showed no effect on FLuc∗ activity when the reporter enzyme was targeted to the cytosol in a reducing compartment (Fig. 8A, right panel). Overall, this finding is consistent with the induction by ritonavir of oxidative stress in the ER promoting the formation of disulfide bonds. This could lead to the misfolding of HBs in ritonavir-treated cells and we thus explored this possibility.

**Fig. 8.**
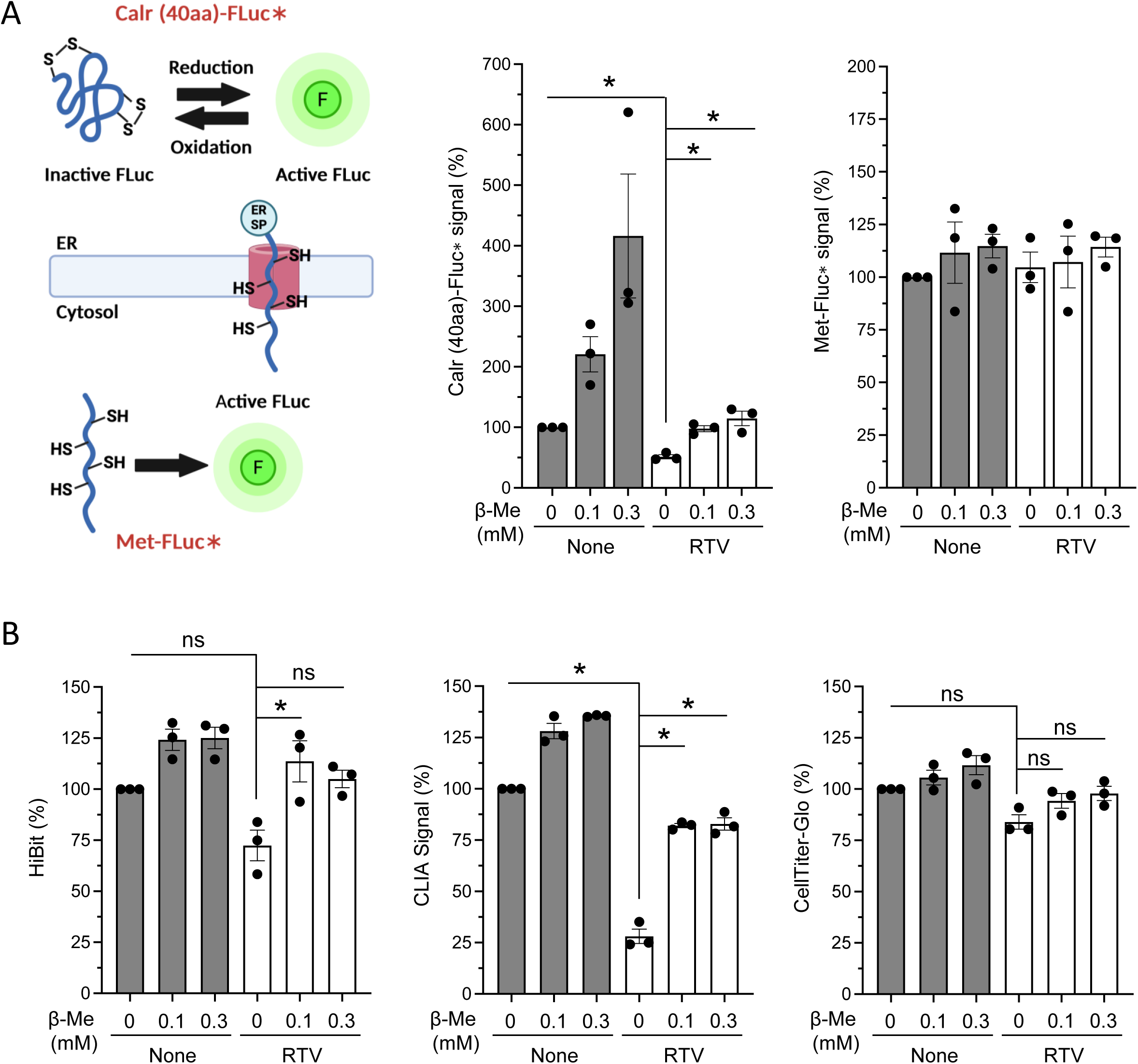
Redox-sensitive luciferase reporter reveals ER hyper-oxidation induced by ritonavir and its impact on S-HBs folding. **(A)** Left panel: Schematic representation of a prospective reporter system to detect disulfide bond formation in the endoplasmic reticulum (ER). The modified firefly luciferase FLuc∗, containing eight additional cysteines, is targeted either to the ER (Calr(40 aa)-FLuc∗) or the cytosol (Met-FLuc∗). Renilla luciferase (RLuc) serves as an internal transfection and expression control. Middle and right panels: HEK293T cells were transfected with Calr(40 aa)-FLuc∗ or Met-FLuc∗ and treated with ritonavir (10CµM), β-mercaptoethanol (β-ME), or both in combination. Luciferase activity was measured to assess the redox status of the ER and the impact of treatments on protein folding. **(B)** Effect of β-ME on the ritonavir-induced impairment of S-HBs parameters after 48 h of treatment in Huh7(S-HBs-HiBiT) cells. Secretion of S-HBs was quantified via the HiBiT signal, proper folding was assessed by CLIA detecting the conformational antigenic loop (HBsAg), and cell viability was measured using CellTiter-Glo assay. All results are expressed as percentages relative to the DMSO-treated control. For all panels, each black dot represents an individual biological replicate. Data correspond to the mean ± SEM of independent experiments (raw values were first normalized to DMSO control in each experiment). One sample t-test using 100% as reference when comparing ritonavir to DMSO, and two-tailed t-test when comparing ritonavir to ritonavir + β-ME. Holm-Šidák correction for multiple testing.

Based on this observation, we asked whether β-ME could revert the effect of ritonavir on S-HBs. To this end, Huh7(S-HBs-HiBiT) cells were treated with ritonavir in the absence or presence of β-ME at 0.1 or 0.3 mM (Fig. 8B). After 48 h of culture, culture supernatants were collected to quantify S-HBs expression using the HiBiT tag and HBsAg level by CLIA. The effect of ritonavir on the HiBiT signal, which reflects the overall secretion of S-HBs, was limited as reported in Fig. 3B, and was reverted by the addition of β-ME. As expected, the CLIA signal was strongly reduced in the presence of ritonavir and importantly, this inhibition was almost fully reverted by β-ME. Finally, cell viability was virtually unaffected by the different treatments. These results demonstrate that β-ME reverts the effect of ritonavir on oxidative stress and the misfolding of S-HBs.

## DISCUSSION

In this study, we have shown that ritonavir alters HBs biogenesis, thus reducing the infectivity of secreted HDV particles. Results support an effect of ritonavir on the folding and the glycosylation pattern of S-HBs, and on the stoichiometry of HBs isoforms. This effect on HBs was also observed with lopinavir and cobicistat, two peptidomimetics structurally related to ritonavir as well as with ketoconazole, an antifungal imidazole. These drugs are all powerful inhibitors of the CYP450 enzymes that metabolize drugs, in particular CYP3A4/5 and, to a lesser extent, others such as CYP2B6, CYP2C9, CYP2C19, and CYP2D6 [39,40]. Transcriptomic and metabolomic analyses provided clear evidence of oxidative stress, ER stress, and activation of the UPR in ritonavir or lopinavir-treated cells as assessed by decreased GSH/GSSG ratio, induction of ophthalmic acid, and increased expression of several genes of the UPR regulon. These observations are consistent with previous reports showing that ritonavir and related HIV protease inhibitors induce such a cellular stress *in vivo* and *in vitro* in different cellular models, including hepatocytes [23–27,41–43]. A molecular probe specifically designed to detect redox changes in the ER provided evidence of increased levels of oxidation in this compartment, assessed by the increased formation of disulfide bonds. Finally, we showed that the reducing agent β-ME could revert the effect of ritonavir on HBsAg expression. Taken together, these observations demonstrate that oxidative stress induced by pharmacological inhibitors of CYP450 enzymes, in particular CYP3A4/5 inhibitors, leads to altered protein folding in the ER and thus, induction of the UPR. In the context of HBV and HDV infections, this has major consequences on the synthesis and functionality of HBV glycoproteins, as these contain multiple disulfide bonds essential to their synthesis and functionality. Overall, our results open perspectives for the design of novel antiviral therapies against HBV and HDV.

A key finding of our study is the alteration of S-HBs folding in cells treated with ritonavir and other CYP3A4/5 inhibitors. These drugs induce an accumulation of the glycosylated form of S-HBs, whereas the non-glycosylated forms are significantly reduced. This abnormal glycosylation profile is associated with a misfolding of the antigenic loop, as assessed by a loss of HBsAg. These two phenotypes were previously reported for mutant of S-HBs where two cysteines involved in disulfide bond formation, C147 and C149, were replaced by serine residues [18,19]. This supports our model where oxidative stress in the ER alters disulfide bond formation, and thus leads to the misfolding of S-HBs. Indeed, upon oxidative stress, disulfide bonds may form too quickly without the possibility for Protein Disulfide Isomerases (PDIs) to reorganize the disulfide bond network. This leads to the misfolding of S-HBs, compromising its structural integrity and function. It has been well established that when conformation of the antigenic loop within HBs is lost, so is infectivity [19,44–46]. Besides, it has been shown that thiol-disulfide exchanges are essential in membrane fusion [19,47]. This reorganization that is essential for virus entry could be impaired by the formation of additional and improper disulfide bonds during the biogenesis of S-HBs. Finally, the larger form of HBs (L-HBs or PreS1) that is essential for entry receptor binding tends to disappear in ritonavir-treated cells as it probably becomes unstable. These alterations in HBs biosynthesis could have profound implications on viral entry as it could impair virus interactions with host cell membranes and fusion mechanisms. This was confirmed by reinfection assays, showing that ritonavir reduces the ability of secreted HDV virions to infect target cells, whereas viral replication was marginally or not affected. Previous studies have also shown that the abundant secretion of HBsAg by infected hepatocytes is a key determinant in the virus ability to evade immune recognition by inducing tolerance [48]. Therefore, conformational changes in S-HBs induced by ritonavir could restore an effective immune response against HBV, and promote immune-mediated clearance of infected cells. Interestingly, two independent case reports describe a complete clearance of HBV in patients treated with high doses of ritonavir [49,50]. These observations suggest that ritonavir’s ability to alter HBs conformation may indeed facilitate HBV suppression *in vivo*.

We then sought to establish a link between CYP3A inhibition and these effects on HBs biogenesis. Major biological roles of CYP450s are the oxidation of xenobiotics to facilitate their secretion, and the biosynthesis of several metabolites such as steroids and vitamins. Although the induction of oxidative stress upon CYP450 inhibition by ritonavir and related molecules is relatively well documented in the literature, molecular mechanisms are not fully understood and several hypotheses can be made. Metabolite monooxygenation by CYP450 depends on NADPH consumption by the flavoprotein POR (CYP450 oxidoreductase), which is followed by the transfer of electrons to CYP450 [51]. The electrons are then used by CYP450s to activate dioxygen and hydroxylate their substrates. However, the degree of coupling between NADPH consumption by POR and substrate oxidation by CYP450s is low, since electron leakage and the formation of reactive oxygen species (ROS) also take place. Such uncoupling has also been reported in particular when a substrate is bound to the CYP450s but is not metabolized [51]. As a consequence, a significant fraction of activated oxygen is released in the form of ROS. Finally, ritonavir and related compounds are ultimately degraded by CYP450s, and thus GSH is consumed by conjugation to ritonavir metabolites in the course of phase II xenobiotic metabolism to facilitate their secretion as glutathione S-conjugates [52]. Although the relative contribution of these mechanisms in the induction of oxidative stress by ritonavir requires further investigations to be determined, it is interesting to notice that POR and CYP450 assemble, with other factors, on the cytosolic face of the ER membrane to form the microsomal monooxygenase (MMO) multienzyme system. Thus, uncoupling of electron transfer by CYP450 inhibition could lead to the production of ROS in the vicinity of the HBsAg production site. The localized diffusion of ROS across the ER membrane could thereby alter disulfide bond formation and protein folding in the ER lumen. Interestingly, S-HBs contains 14 cysteine residues for a total of 226 amino acids (*i.e.*: 6.2%), which potentially makes it particularly sensitive to abnormally high oxidative conditions in this organelle.

Protein misfolding in the ER upon ritonavir treatment was assessed by the expression of key UPR markers such as CHOP. Several enzymes involved in the serine-glycine pathway and folic acid cycle also had their expression induced (Fig. 5D). These enzymes were previously associated with the UPR regulon and their induction leads to increased metabolic flux for glycine production [31]. Our transcriptomic data also showed the induction of CTH (Cystathionine gamma-lyase), an enzyme of the transsulfuration pathway which is key for cysteine synthesis from methionine and serine. Both glycine and cysteine are essential for GSH synthesis which, as a reducing agent, helps to control oxidative stress in general. More specifically, GSH can be imported into the lumen of the ER via the Sec61 translocon to regulate redox balance for controlling disulfide bond formation [53]. We also observed an accumulation of tripeptide ophthalmic acid in ritonavir-treated cells, a phenomenon also reported in animals treated with this drug [54]. This tripeptide metabolite is also induced by acetaminophen, the prototype of hepatotoxic drug whose byproduct, N-acetyl-p-benzoquinone imine (NAPQI), induces an oxidative stress by depleting cells in GSH [36]. Ophthalmic acid is produced by the same enzymes than GSH, *i.e.:* Glutamyl-Cysteine Ligase (GCL) and Glutathione Synthetase (GSS), but using 2-aminobutyric acid instead of cysteine [36,37]. This metabolite is synthetized by transamination of 2-oxobutyric acid, a product of cystathionine cleavage by CTH to produce cysteine. Increased level of ophthalmic acid reflects the induction of GSH synthesis in ritonavir-treated cells [36], and is a marker of oxidative stress that corroborates the GSH/GSSG ratio and the low signal obtained with the ER-specific probe FLuc*. It is also interesting to note that a recent report has challenged the accepted model about serine/glycine homeostasis by showing that, under physiological conditions in mice, glycine is metabolized into serine in the liver and not the opposite [55]. In the future, it would be interesting to determine whether ritonavir administration can influence this balance *in vivo* by reverting this flux, promoting serine conversion to glycine as suggested by our *in vitro* observations.

In conclusion, this study highlights the pivotal role of cytochrome P450 enzymes in regulating the structure and glycosylation of S-HBs. Through the use of various hepatic cell lines, including dHepaRG, PHH and engineered Huh7-derived cells, we demonstrate that cytochrome P450 inhibition significantly alters the viral envelope proteins, resulting in the reduced infectivity of HDV particles. Our findings suggest that targeting cytochrome P450 enzymes may offer a novel therapeutic approach for modulating HBV and HDV infections. The ability to manipulate viral protein maturation and infectivity through metabolic pathways opens new avenues for antiviral drug development, particularly in the context of chronic viral infections.

## ACKNOWLEDGEMENTS

The project was funded by an intramural CIRI grant (AO-6-2020), the SATT Pulsalys (Project Delta-i), INSERM-Transfert (Project IPP-Houra) and the ANRS - Maladies infectieuses émergentes (ECTZ208004). WEO was funded for his Ph.D by the ANRS - Maladies infectieuses émergentes (ECTZ208715). We thank the Central Institute for Experimental Medicine and Life Science (Kawasaki, Japan) for providing HepaSH®.

---

**Figure S1.**
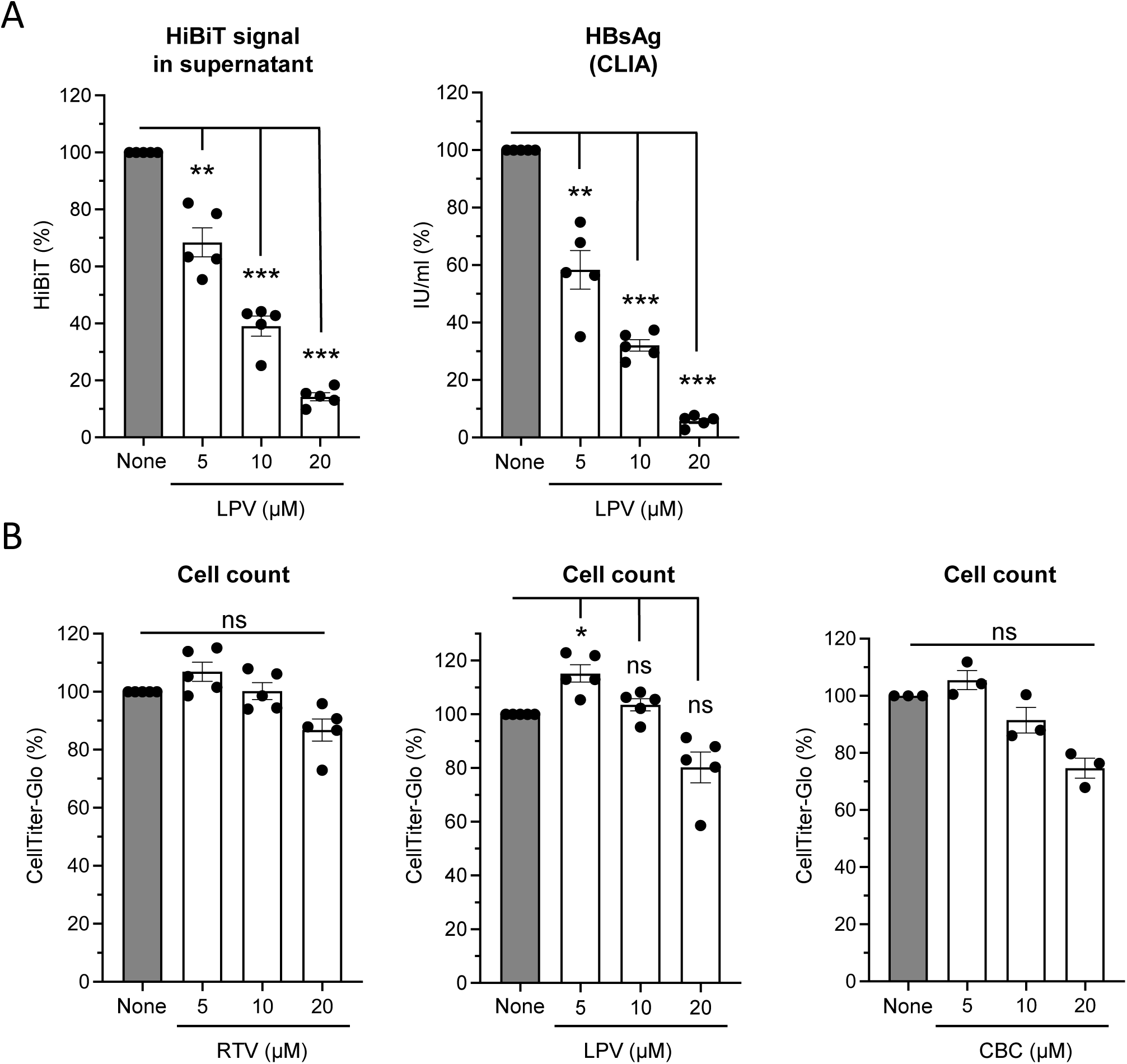

**Figure S2.**
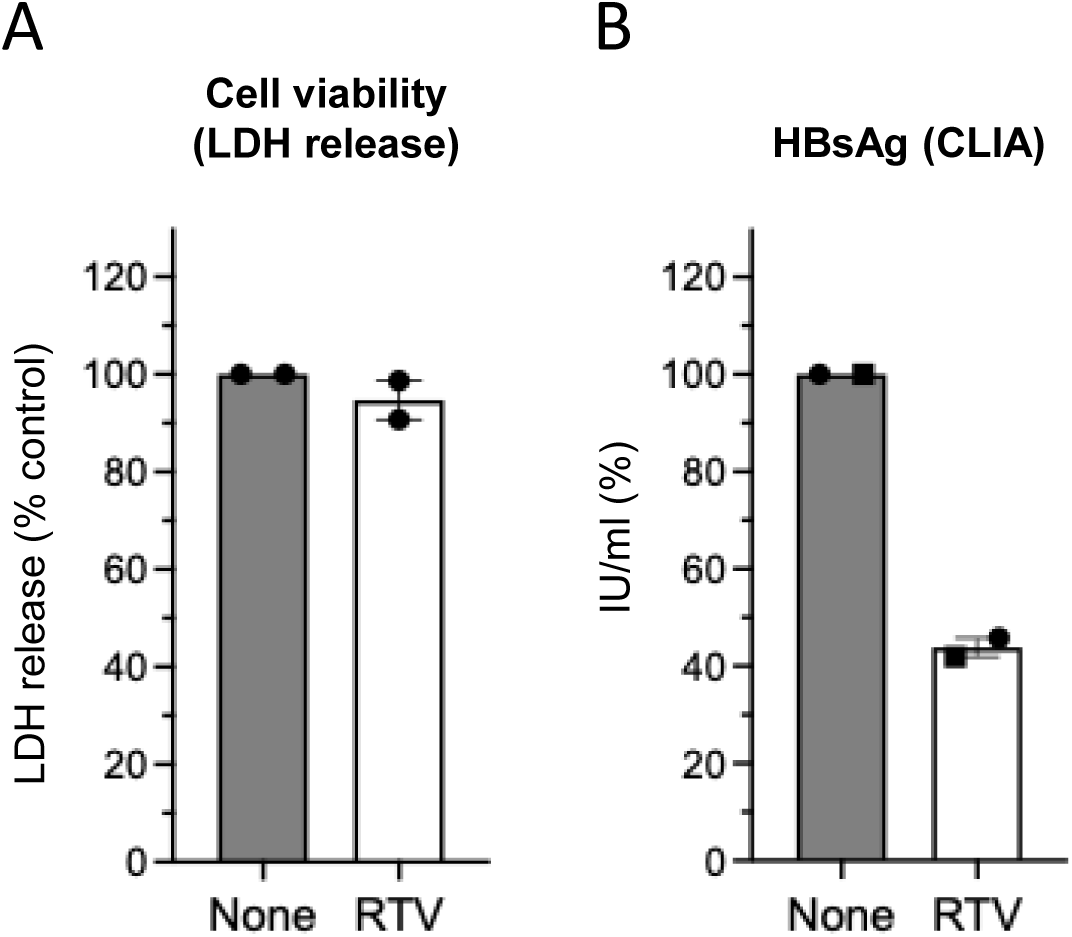

**Figure S3.**
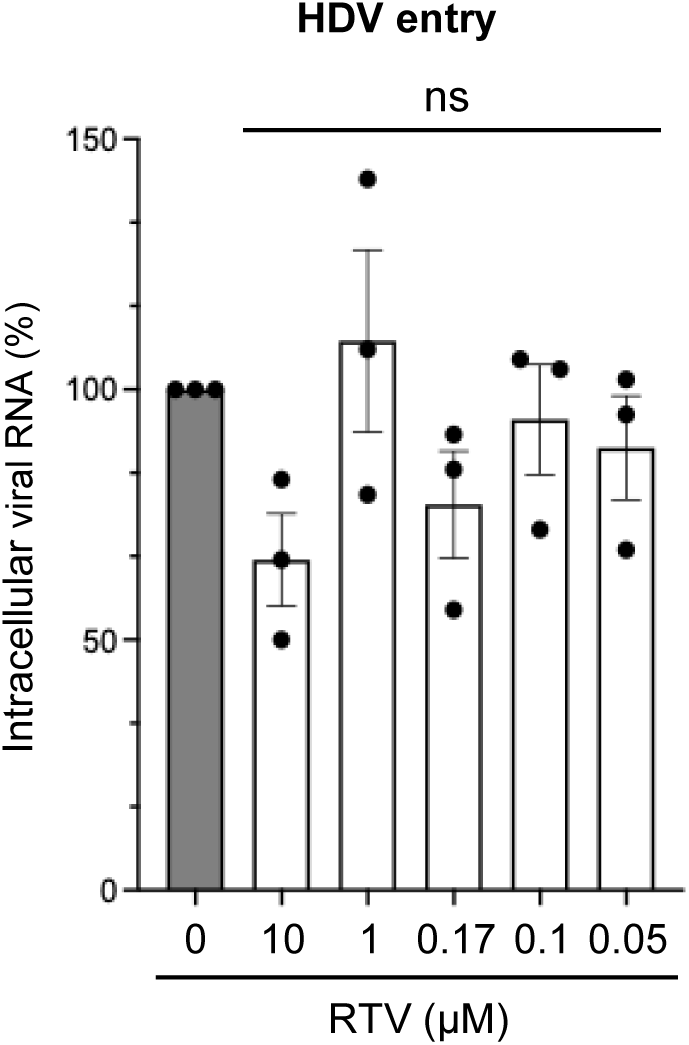

**Figure S4.**
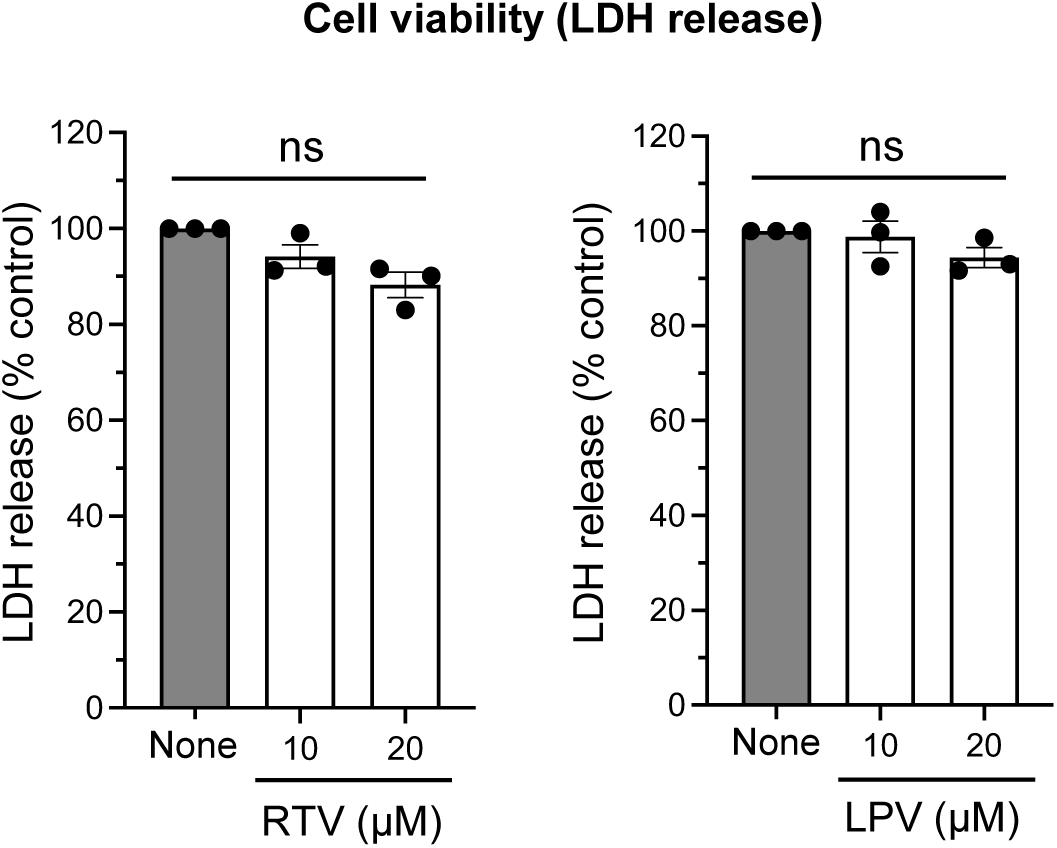

**Figure S5.**
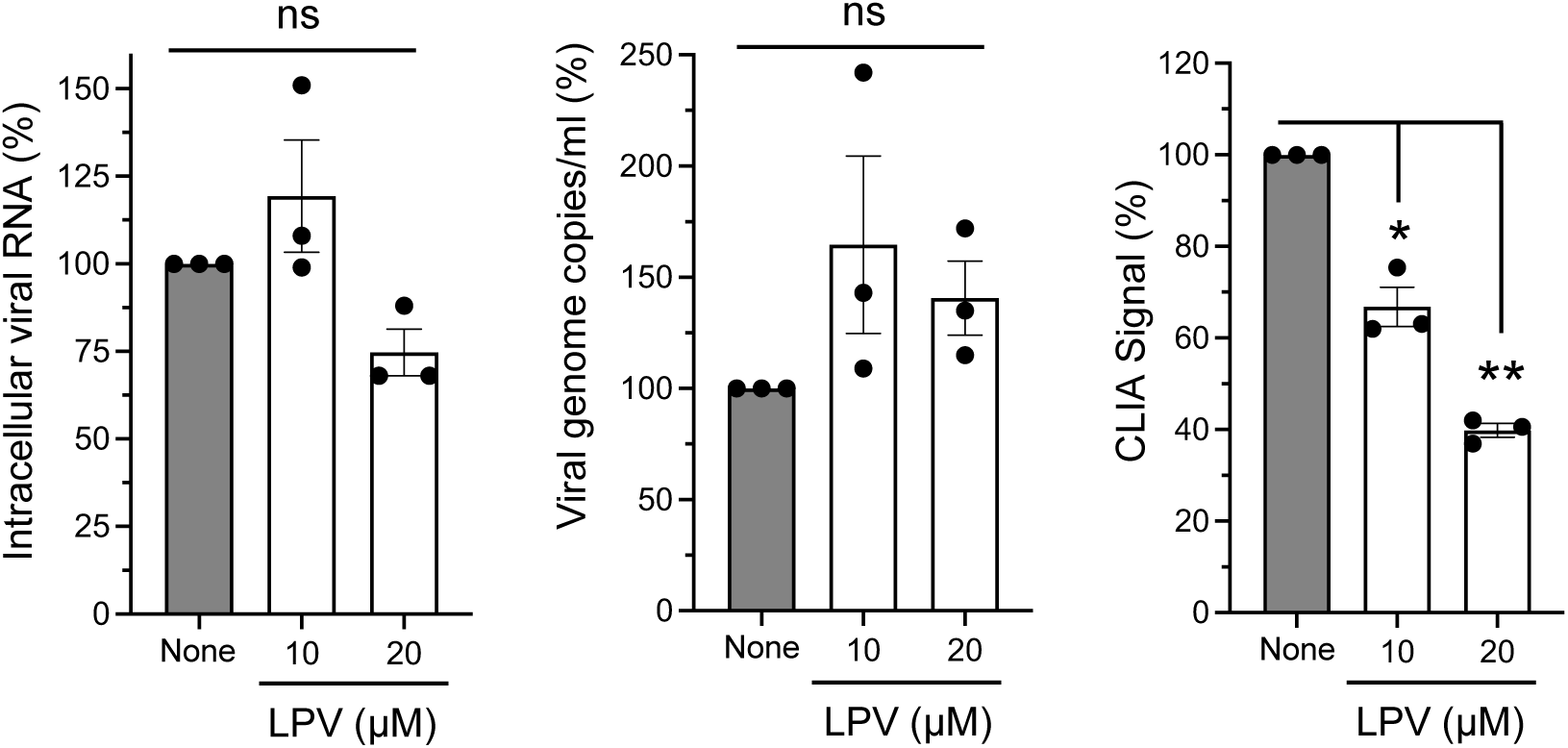

**Figure S6.**
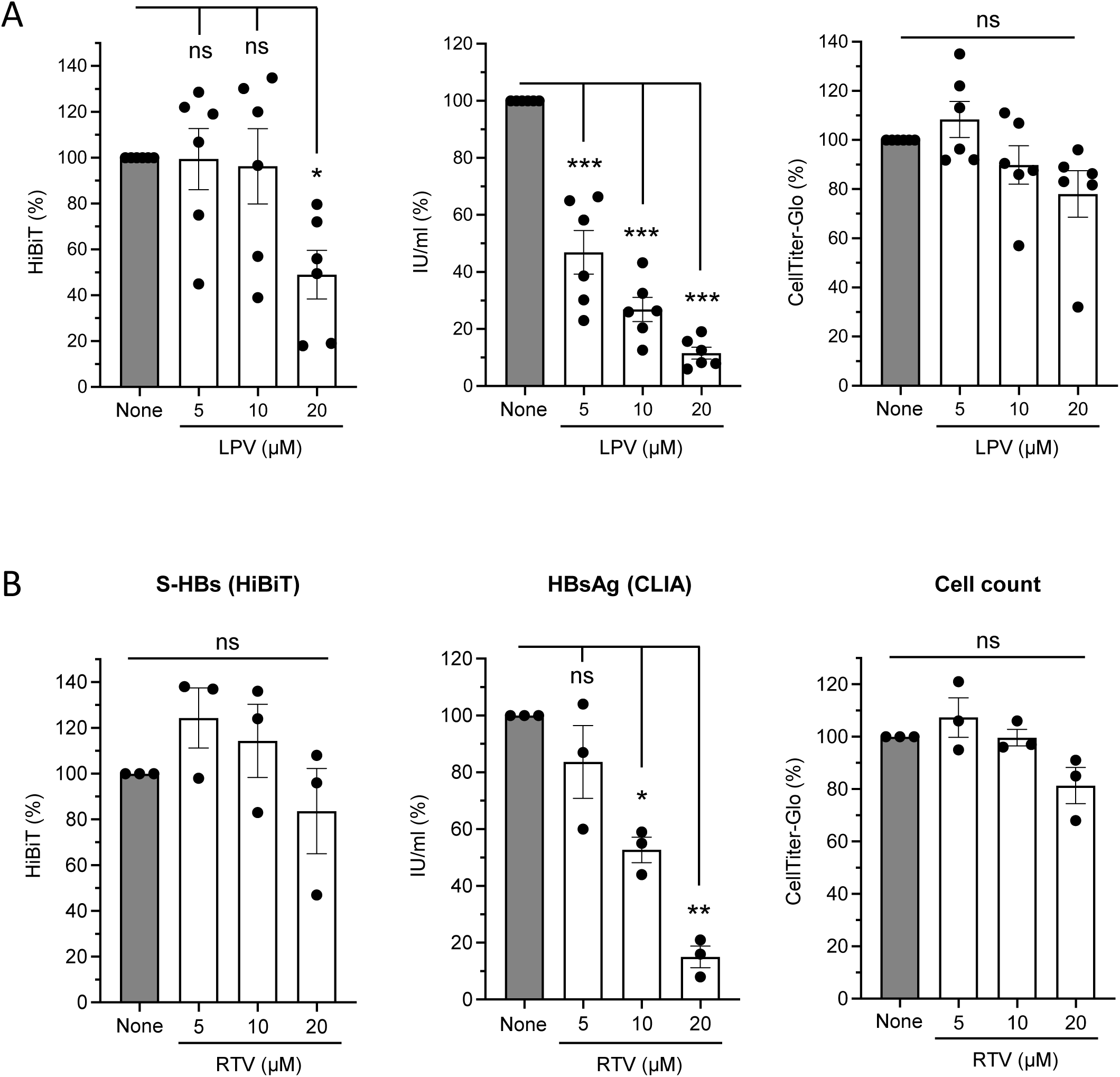

**Figure S7.**
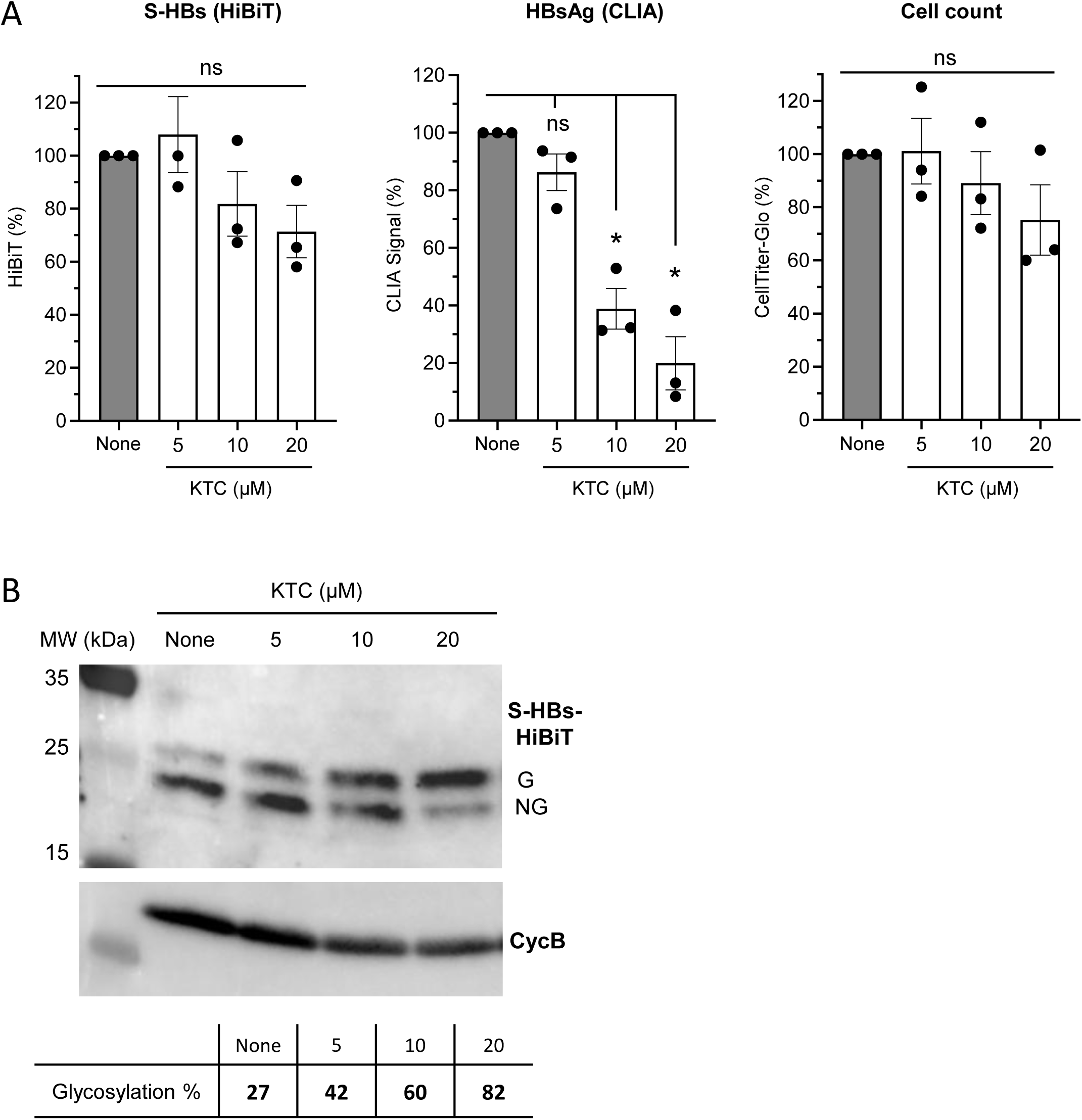

**Figure S8.**
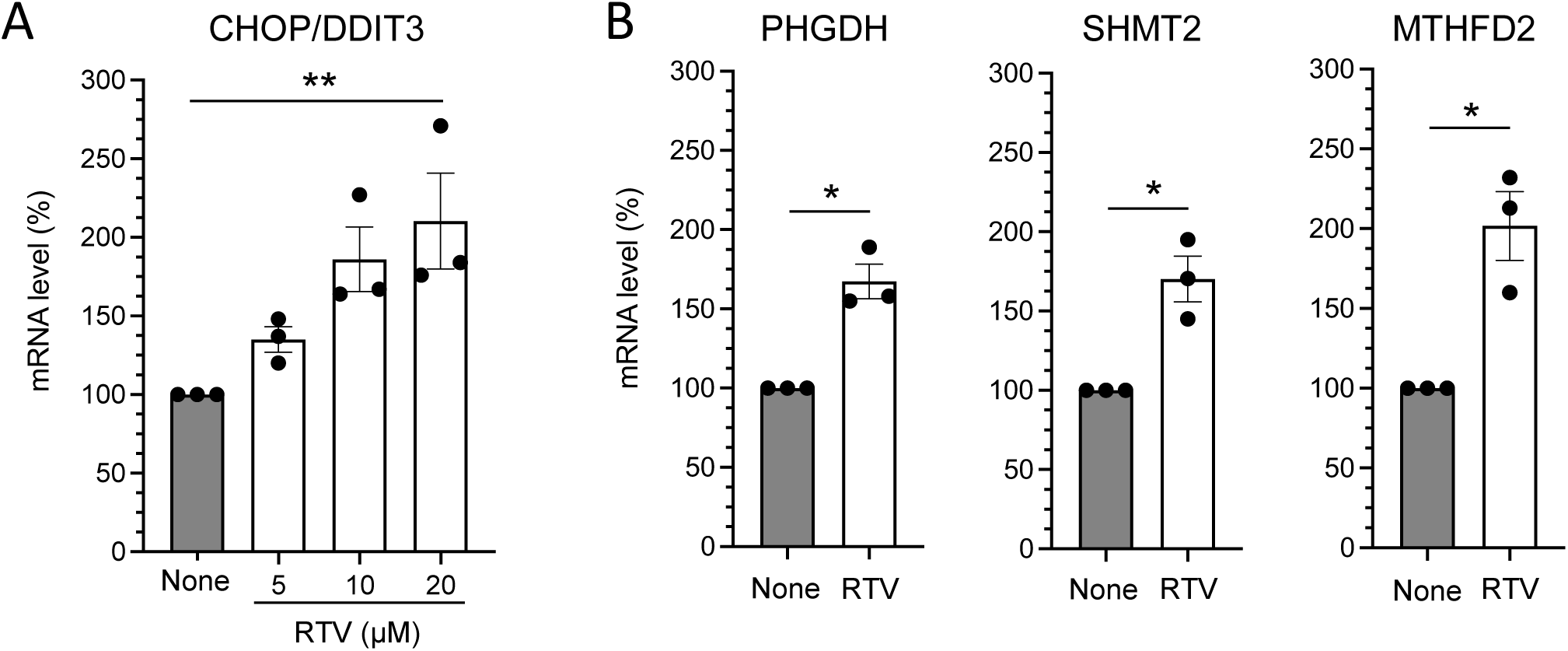

