## Supplementary Data for "Ritonavir-Induced Cellular Stress Alters Viral HBs Glycoprotein Biogenesis and Production of Infectious Hepatitis D Virions"

* Equally contributors to this work; ^$^ Co-supervision of this project; ^1^ Equipe VIRIMI, CIRI, Centre International de Recherche en Infectiologie, Univ. Lyon, Inserm U1111, CNRS UMR5308, ENS-Lyon, Université Claude Bernard Lyon 1, France ; ^2^ Equipe Hepvir, CIRI, Centre International de Recherche en Infectiologie, Univ. Lyon, Inserm U1111, CNRS UMR5308, ENS-Lyon, Université Claude Bernard Lyon 1, France ; ^3^ Laboratoire de Biologie Tissulaire et d'Ingénierie Thérapeutique, Univ Lyon, CNRS, Université Claude Bernard Lyon 1, UMR 5305, 7 Passage du Vercors, CEDEX 07, 69367 Lyon, France ; ^4^ Cell Biology Center, Institute of Integrated Research, Institute of Science Tokyo, Yokohama, Kanagawa 226-8501, Japan ; ^5^ Chemical Proteomics, Dept. of Medical Biochemistry and Biophysics (MBB), Biomedicum A9, Karolinska Institutet, SE 17177 Stockholm, Sweden ; ^6^ Labéo, EA 7450 Biotargen, Normandie Université, UniCaen, 14053 Caen, France; ^7^ INSERM U1259, Université de Tours, 37000 Tours, France ; ^8^ Service de Virologie, Institut des Agents Infectieux, Hôpital de la Croix Rousse, Hospices Civils de Lyon, Lyon, France ; ^9^ Structure et Instabilité des Génomes (StrInG), Muséum National d'Histoire Naturelle, INSERM U1154, CNRS UMR 7196, Alliance Sorbonne Université, 75005 Paris, France. ; ^10^ Laboratoire P4 INSERM-Jean Mérieux, 69007 Lyon, France.

**TABLE OF CONTENT**

- **Material and methods**
- **Supplementary Figures 1-8 and legends**
- **References**

**MATERIAL AND METHODS**

**Cells and viruses**

Huh7 cells (kindly provided by Dr. Marco Binder; Heidelberg University; Germany) and HEK-293T cells (CRL-3216, ATCC) were cultured at 37°C and 5% CO_2_ in high-glucose DMEM (4.5 g/L glucose; Ref. 10566016; Gibco, France) supplemented with 10% fetal calf serum (FCS; Corning, Life Sciences) and 100 U/mL penicillin/streptomycin (Gibco, France). HBV viral stocks (Genotype D) were obtained by collecting supernatants from HepAD38 cells as previously described [1]. HDV stocks were produced by co-transfecting Huh7 cells with the pT7HB2.7 plasmid encoding HBV envelope proteins (Genotype D) and the pSVLD3 plasmid, which contains a head-to-tail trimer of the full-length HDV cDNA (Genotype 1), allowing transcription of genomic HDV RNA from the simian virus 40 late promoter [2]. Primary Human Hepatocytes (PHH) were prepared from a human liver resection obtained from the Centre Léon Bérard (Lyon) with French ministerial authorizations (AC 2013-1871, DC 2013–1870, AFNOR NF 96 900 September 2011) as previously described [3]. HepaSH®, which correspond to PHH amplified in TK-NOG mice with a humanized liver, were purchased from Wepredic (Saint-Grégoire, France) and provided by the Central Institute for Experimental Medicine and Life Science (Kawasaki, Japan)[4]. PHH and HepaSH® were cultured in William’s E medium (Ref. 12551032; Gibco, France) supplemented with 5% FCS (HyClone™ FetalClone™ II, Cytiva), 100 U/mL penicillin/streptomycin (Gibco,France), 5 μg/mL insulin (Sigma-Aldrich), 50 μM hydrocortisone hemisuccinate (Sigma-Aldrich), 1x non-essential amino acid solution (Gibco), 2 mM L-glutamine (Gibco), EGF (2 ng/ml), 2% DMSO, 10 mM HEPES (Gibco). HepaRG cells were cultured in William’s E medium (Ref. 12551032; Gibco, France) supplemented with 10% FCS (HyClone™ FetalClone™ II, Cytiva), 100 U/mL penicillin/streptomycin (Gibco,France), 5 μg/mL insulin (Sigma-Aldrich), and 50 μM hydrocortisone hemisuccinate (Sigma-Aldrich). To induce differentiation, cells were cultured for two weeks in 24-well plates, and for two additional weeks in culture medium supplemented with 1.8% DMSO. Cells were infected overnight in the presence of 4% PEG using either HBV, HBV and HDV, or PHH supernatants containing infectious virions as previously described [5,6]. After 24 h, cells were washed twice with PBS to remove the inoculum and fresh medium was added to the wells. All cell lines were regularly tested for mycoplasma contamination using Mycoplasmacheck (Eurofins Genomics, Germany).

**Reagents**

Ritonavir, lopinavir, cobicistat, and ketoconazole were purchased from MedChemExpress. Lonafarnib was purchased from Sigma-Aldrich. Stock solutions were prepared at 10 mM in DMSO, stored at -80°C, and used at the indicated concentrations. β-mercaptoethanol was provided as a ready-to-use solution for cell culture at a concentration of 50 mM (Thermo Fisher).

**Generation of Huh7(HiBiT-HDAg-L/S-HBs) and Huh7(S-HBs-HiBiT) reporter cells**

The HiBiT tag corresponds to an 11-amino-acid fragment of NanoLuc luciferase, which spontaneously reconstitutes a functional bioluminescent enzyme when the complementary NanoLuc fragment is added extemporaneously. The genes encoding HiBiT-HDAg-L and S-HBs-HiBiT were obtained by chemical synthesis and cloned into pDONR221 (Invitrogen). The HiBiT tag was separated by the AGSSGGSSGA linker followed by the HDAg-L sequence (AJ000558.1; Genotype 1). The S-HBs sequence (EU594400; Genotype D2) was separated by the AGSSGGSSGA linker followed by the HiBiT tag. The sequences encoding HiBiT-HDAg-L and S-HBs-HiBiT were transferred by Gateway cloning from pDONR221 into pLEX_307 (a gift from David Root; Addgene #41392; Addgene, Watertown, MA, USA) to produce lentiviral particles. Huh7 cells were transduced and selected for 10 days in culture medium with puromycin (1 μg/mL) to obtain the Huh7(HiBiT-HDAg-L) cell line and the Huh7(S-HBs-HiBiT) cell line expressing HiBiT-HDAg-L and S-HBs-HiBiT, respectively. To demonstrate that the HiBiT-HDAg-L protein is secreted when S-HBs is co-expressed, Huh7(HiBiT-HDAg-L) cells were transfected with pCI-neo-HBs-S, a plasmid expressing S-HBs. To generate this plasmid, the S-HBs sequence was amplified by PCR from pT7HB2.7 (Genbank ID: V01460.1; HBV strain ayw), cloned into pDONR223, and then transferred by Gateway recombination into a pCI-neo expression vector (Promega) modified for being compatible with this cloning strategy (kindly provided by Dr. Yves Jacob). Huh7(HiBiT-HDAg-L) cells were plated in 24-well plates (5x10^4^ cells/well), and then transfected with 500 ng of plasmid using jetOPTIMUS DNA transfection reagent following manufacturer’s recommendations (Polyplus, France). Supernatants were collected and analyzed 48 h later. To generate a cell line stably expressing S-HBs, Huh7(HiBiT-HDAg-L) cells were transduced with a lentiviral vector. The S-HBs sequence was first transferred by Gateway recombination from pDONR223 into the pLenti-PGK-Neo-DEST vector to produce lentiviral particles. Then, Huh7(HiBiT-HDAg-L) cells were transduced with the corresponding particles, and selected for ten days in culture medium supplemented with both G418 (1 mg/mL) and puromycin (1 μg/mL) to obtain the Huh7(HiBiT-HDAg-L/S-HBs) cell line.

**Detection of secreted HiBiT-HDAg-L or S-HBs-HiBiT**

To quantify the secretion of HiBiT-HDAg-L or S-HBs-HiBiT, Huh7 cells stably the corresponding constructs were seeded in 96-well plates at 7x10^3^ cells/well in 200 µL of culture medium. One day later, 100 µL of supernatant was replaced with fresh medium containing either the drug of interest or DMSO alone. After 48 h of incubation, culture supernatants were collected and centrifuged for 5 min at 1200 rpm. Then, 50 µL of the supernatant were transferred into white-bottom 96-well cell culture plates, and 50 µL of NanoGlo HiBiT Lytic Reagent (Promega, USA; Ref. N3030), a buffer containing the recombinant N-terminus of Nano luciferase (LgBiT) and the Nano luciferase substrate Furimazine, were added. Luminescence intensity was then measured using a microplate reader (Tristar 5, Berthold, Germany). For measurements in the cell lysate, 50 µL of the medium were left in contact with the cells, and 50 µL of NanoGlo HiBiT Lytic Reagent was added to lyse the cells. The subsequent steps were identical to those described above.

**HBs antigen quantification (CLIA)**

The levels of secreted HBsAg in cell culture supernatants were quantified using an ELISA-based chemiluminescence immunoassay kit (Autobio, China), following the manufacturer's instructions.

**Quantification of cell proliferation**

Cell counts in culture wells were determined using the CellTiter-Glo Luminescent Cell Viability Assay (G7570; Promega, France), which quantifies ATP as a proxy for the number of viable cells. Cells were incubated for 48 h in white 96-well plates with 200 µL of culture medium containing either DMSO alone or the indicated drugs. For ATP quantification, 150 µL of culture medium were first removed from each well before adding 50 µL of CellTiter-Glo reagent. After mixing and 10 min of incubation at room temperature (RT), luminescence was measured using a microplate reader (Tristar 5, Berthold, Germany).

**Screening of the APExBio chemical library**

The APExBio chemical library (Metabolism-related Compounds; DiscoveryProbe; Ref. L1032; APExBio Technology) was purchased from Stratech. The 493 drugs of the library were provided in solution at 10 mM in DMSO unless specified otherwise by the manufacturer. The library was stored at −80°C. For the screening, white 96-well plates (Greiner) containing the drugs were seeded with 1x10^4^ Huh7(HiBiT-HDAg-L/S-HBs) cells per well. Each drug was screened at 10 μM in a final volume of 200 μL. After 48 h of culture, supernatants were collected to quantify, as described above, the levels of secreted HiBiT-HDAg-L and HBsAg using the NanoGlo HiBiT Lytic Reagent (Promega) and the CLIA (Autobio), respectively. Cell viability in presence of tested drugs was assessed using the CellTiter-Glo luminescent assay. Each screening plate contained four positive and four negative control wells treated with lonafarnib (0.2 μM) and DMSO, respectively. For each readout (HiBiT, CLIA, CellTiter-Glo), raw signals of the plate were normalized to the mean of the signal obtained with the four DMSO control wells and were expressed as percentages.

**LDH assay**

The LDH-Glo Cytotoxicity Assay (Promega) was used following manufacturer’s recommendations to measure the viability of infected PHH or dHepaRG cells treated with DMSO, ritonavir or lopinavir. Supernatants of PHH or dHepaRG cells were diluted 100-fold by mixing 2 µL of supernatant with 198 µL of LDH Storage Buffer (200 mM Tris-HCl, pH 7.3, 10% glycerol, 1% BSA). Then, 50 µL of the diluted sample were mixed with 50 µL of LDH Detection Reagent (LDH Detection Enzyme Mix + Reductase Substrate) in a 96-well plate. After 45 min of incubation at RT, luminescence was measured using the Tristar 5 Multimode Reader (Berthold, Germany). For HepaRG cells, results were expressed as percentage of viability using untreated cells and lysed cells (Triton X-100 10%) as negative and positive controls, respectively. For PHH, LDH release was expressed as percentage relative to DMSO-treated control wells.

**Real Time Quantitative PCR**

For all intracellular quantifications (endogenous genes or viral RNA), cells were washed with PBS and processed using the Monarch Total RNA Miniprep Kit (New England BioLabs, France). After RNA quantification, reverse transcription was performed using the Maxima First Strand cDNA Synthesis Kit for RT-qPCR (Thermo Fisher Scientific, #K1641), following the manufacturer’s recommendations. The qPCR reactions were carried out using the PowerTrack SYBR Green Master Mix (A46012; Applied Biosystems) on a CFX96 Real-Time PCR Detection System (Bio-Rad, France). Gene induction was expressed as fold change using the 2^−ΔΔCt^ method, with human RPL13A and PrP as housekeeping genes. To quantify HBV and HDV viral genomes in the culture supernatant, 150 μL of supernatant was collected after homogenization and mixed with 600 μL of RAV1 lysis buffer (Macherey-Nagel Virus Kit). The mixture was then incubated at 70°C for 5 min. Following this step, nucleic acids were extracted according to the manufacturer’s instructions to isolate either RNA (for HDV) or DNA (for HBV). For HDV quantification, the Luna Universal Probe One-Step RT-qPCR Kit (New England Biolabs) was used with a Taqman probe (FAM-CTCTTCTTCCTCCTTGCTGA-BHQ1)[7]. HBV viremia was determined by quantification of viral genomes in supernatants using qPCR and the SYBR Green Master Mix (A46012; Applied Biosystems).

*Table 1: Primers used for RT-qPCR*

| **Name** | **Forward Primer** | **Reverse Primer** |
| --- | --- | --- |
| HBV | ACCGAATGTTGCCCAAGGTC | TATGCCTCAAGGTCGGTCGT |
| HBV viremia | GGAGGGATACATAGAGGTTCCTTGA | GTTGCCCGTTTGTCCTCTAATTC |
| HDV | CGGGCCGGCTACTCTTCT | AAGGAAGGCCCTCGAGAACA |
| SHMT2 | GCTCAACCTGGCACTGACTG | CACTGATGTGGGCCATGTCT |
| PHGDH | CTTACCAGTGCCTTCTCTCCAC | GCTTAGGCAGTTCCCAGCATTC |
| MTHFD2 | GCAGGAGGTAGAAGAGTCGGT | TCTGGAAGAGGCAACTGAACAA |
| CHOP/DDIT3 | AAGGCACTGAGCGTATCATGT | TGAAGATACACTTCCTTCTTGAAC |
| PrP | TGCTGGGAAGTGCCATGAG | CGGTGCATGTTTTCACGATAGTA |
| RPL13A | AAAAGCGGATGGTGGTTCCT | GCTGTCACTGCCTGGTACTT |

**Western-blot analyses**

Cell lysates from 3x10^6^ cells were prepared in 1X RIPA lysis buffer (MERCK, 20-188) supplemented with 1% protease inhibitor cocktail (Sigma-Aldrich, P8340). After the removal of insoluble material by centrifugation at 13,000 rpm, clarified protein lysates were collected and stored at -80°C. Total proteins were quantified (DC Protein Assay; Bio-Rad; France), separated by SDS-PAGE, and analyzed by Western blot on a PVDF membrane. The PVDF membrane was saturated in blocking buffer (TBST supplemented with 0.1% Tween-20 and 5% non-fat milk powder), and the blots were then incubated overnight at 4°C with a rabbit polyclonal HBsR247 primary antibody (a kind gift from Camille Sureau; INTS) in the same buffer (1:1,000 dilution). After washing, incubation with the secondary antibody was performed for 30 min at room temperature. HRP-conjugated anti-rabbit antibody (Cell Signaling Technology, 7074) was diluted 1:10,000 and detected by enhanced chemiluminescence reagents according to the manufacturer’s instructions (SuperSignal Chemiluminescent Substrate, Thermo Fisher Scientific). For the loading control, a rabbit anti-Cyclophilin B antibody (Cell Signaling Technology, 43603) was used at a 1:1,000 dilution. To detect the HiBiT tag, the proteins were prepared as described above. After transfer, the membrane was briefly washed in TBST-0.1% Tween, then incubated for 1 h at room temperature in the LgBiT/buffer solution (Nano-Glo® HiBiT Blotting System; Promega; Ref. N2410). Nano-Glo® Luciferase Assay substrate was added and incubated for 5 min before exposure (ChemiDoc MP Imaging System).

**Transcriptomic analysis**

Three biological replicates were prepared for each condition. Huh7 cells were seeded in a 6-well plate (2 × 10⁵ cells/well) and cultured for 24 h in 3 mL of culture medium. The medium was then removed and replaced with fresh medium supplemented with the appropriate compounds: either DMSO alone, Ritonavir (10 µM), or Lopinavir (10 µM). After 48 h of treatment, the medium was discarded and total RNA was extracted using the Direct-zol DNA/RNA Miniprep kit, following the manufacturer's instructions (Zymo Research). RNA samples were stored at -80 °C prior to shipment on dry ice. DNA library construction, sequencing and raw data analysis described in these sections were performed by Microsynth AG (Balgach, Switzerland). Total RNA was quantified using Quant-iT RiboGreen (Thermofisher). Integrity was checked on Fragment analyzer (Agilent) with a kit suitable for RNA. Libraries were prepared using the Illumina® Stranded mRNA Prep, Ligation (96 Samples) (20040534) and 300 ng total RNA input. Final libraries were quantified using pico488 (Lumiprobe) and library size was checked on Fragment analyzer (Agilent). Equimolar amounts of the samples were pooled prior to sequencing. Subsequently the Illumina NovaSeq platform and an SP, 300 cycles kit were used to sequence the libraries in paired-end fashion. The produced paired-end reads which passed Illumina’s chastity filter were subject to de-multiplexing and trimming of Illumina adaptor residuals using Illumina’s bcl2fastq software version 2.20.0.422 (no further refinement or selection). Quality of the reads in fastq format was checked with the software FastQC (version 0.11.8; https://www.bioinformatics.babraham.ac.uk/projects/fastqc/). Raw reads having average Q-values below 24 or incorporating uncalled ‘N’ bases or being smaller than 25 bases after trimming were filtered using the BBTools software suite (version 38.86; https://sourceforge.net/projects/bbmap/). The splice aware RNA mapping software STAR (version 2.7.10a [8]) was used to map the surviving reads to the *Homo sapiens* ENSEMBL reference genome provided by Illumina (downloaded from iGenomes 12.02.2024). To count the uniquely mapped reads to annotated genes, the software htseq-count (HTSeq version 0.13.5) was used [9]. Normalization of the raw counts and differential gene expression analysis was carried out with help of the R software package DESeq2 (version 1.26.0) [10].

**Metabolomic analysis**

Four biological replicates were prepared for each culture condition. Huh7 cells were seeded in 10 mm dishes (2 × 10⁶ cells/dish) and grown for 24 h in 10 mL of culture medium. The supernatant was removed and replaced with 8 mL of fresh culture medium containing either DMSO alone or Ritonavir (10 µM). After 48 h of culture, the medium was removed and the cells were washed with PBS buffer (37°C) for 60 seconds. The wash solution was discarded and replaced with 5 mM HEPES (37°C) for a second wash of 5 s, then rapidly removed. Metabolites were immediately extracted in 1 mL of ice-cold (-20°C) 80% MS-grade MeOH (900688; Sigma-Aldrich). Samples were then stored at -80°C before shipping on dry ice. Sample analysis was performed by MS-Omics (Denmark) as described below. Semi-polar and polar metabolite profiling was performed by MS-Omics (Vedbæk, Denmark).

The semi-polar metabolite analysis was carried out using a Vanquish LC (Thermo Fisher Scientific) coupled to an Orbitrap Exploris 240 MS (Thermo Fisher Scientific). The UHPLC used an adapted method described by Doneanu *et al.* (UPLC/MS Monitoring of Water-Soluble Vitamin Bs in Cell Culture Media in Minutes, Water Application note 2011, 720004042en). An electrospray ionization interface was used as ionization source. Analysis was performed in positive and negative ionization mode under polarity switching. Untargeted data processing Metabolomics processing was performed untargeted using Compound Discoverer 3.3 (Thermo Fisher Scientific) and Skyline (24.1, MacCoss Lab Software; [11]) for peak picking and feature grouping, followed by an in-house annotation and curation pipeline written in MatLab (2022b, MathWorks). Identification of compounds were performed at four levels; Level 1: identification by retention times (compared against in-house authentic standards), accurate mass (with an accepted deviation of 3ppm), and MS/MS spectra, Level 2a: identification by retention times (compared against in-house authentic standards), accurate mass (with an accepted deviation of 3ppm). Level 2b: identification by accurate mass (with an accepted deviation of 3ppm), and MS/MS spectra, Level 3: identification by accurate mass alone (with an accepted deviation of 3ppm). Annotations on level 2b are based on accurate mass and MS/MS spectra measured with high resolution Orbitrap ESI-MS in mzCloud (Thermo Fisher Scientific), MassBank of North America (UC Davis) and the European MassBank (Helmholtz Centre for Environmental Research Leipzig). The annotations on level 3 are based on searches in the Human metabolome database (version 5.0).

For the polar metabolite analysis, the high-performance liquid chromatography (HPLC) coupled with high-resolution mass spectrometry (HRMS) analysis was performed using a Q Exactive mass spectrometer (Thermo Fisher Scientific) equipped with an electrospray ionization (ESI) source. Chromatographic separation was achieved using a hydrophilic interaction liquid chromatography (HILIC) method optimized for the analysis of polar metabolites. HPLC separation on a ACQUITY Premier BEH Amide column (1.7 µm, 2.1 mm X 100 mm, Waters) employed a two-step gradient elution over 12 minutes with a flow rate of 450 µL/min followed by 4 minutes re-equilibration. Column temperature was maintained at 30°C with an injection volume of 2 µL. Eluent A consisted of 10 mM ammonium formate in ACN, pH 10, and eluent B consisted of 10 mM ammonium formate in 40% ACN, pH 10 with 0.1% medronic acid. Analysis was performed in both positive and negative ionization modes. The Q Exactive mass spectrometer operated at a resolution of 120,000 in a scan range of 60 to 900 m/z. Iterative data-dependent MS/MS (dd-MS²) acquisition was achieved on a TopN of 10 with stepped collision energy (NCE) of 20, 40, 60. System calibration was performed weekly ensuring a mass accuracy below 1 ppm. Annotations on level 2b are based on accurate mass and MS/MS spectra measured with high resolution Orbitrap ESI-MS in mzCloud (ThermoFisher Scientific), MassBank of North America (UC Davis) and the European MassBank (Helmholtz Centre for Environmental Research Leipzig). The annotations on level 3 are based on searches in the Human metabolome database (version 5.0).

**GSH and GSSG measurement**

Huh7(S-HB-HiBiT) cells were seeded in 96-well plates at 7x10^3^ cells/well, and one day later were treated with DMSO or the drugs of interest. After 48 h, cellular GSH and GSSG were measured using the GSH/GSSG-Glo™ Assay Kit (Promega, V6611) following manufacturer’s recommendations.

**Luciferase probe redox status in the ER**

Firefly luciferase (FLuc)-based reporter plasmids pHK1608 (encoding Met-FLuc*) and pSY107 [Calr (40 aa)-FLuc*] were provided by Hiroshi Kadokura (Institute of Science Tokyo). Details of these plasmid constructions are described in reference [12]. Cells were seeded at a density of 1.2x10^5^ cells per well. Twenty-four hours later, cells were transfected with 90 ng of the plasmid of interest (pHK1608 or pSY107) and 10 ng of pTK-Renilla (Promega) for normalization purposes. Six hours post-transfection, the medium was removed and replaced with the corresponding treatment conditions. Forty-eight hours after transfection, the medium was discarded and the cell monolayer was washed twice with PBS before being lysed in 100 µL of in-house lysis buffer for 30 minutes at room temperature under agitation. The lysis buffer was composed of PBS supplemented with 1% NP-40 (Sigma-Aldrich), 1 mM PMSF (MedChemExpress), 1 mM benzamidine (MedChemExpress), and 1 µg/mL pepstatin (MedChemExpress). Ten microliters of the resulting lysate were used to measure Firefly luciferase activity by adding 50 µL of LAR II reagent (Dual-Luciferase® Reporter Assay System, Promega). Subsequently, 50 µL of Stop & Glo® Reagent were added to measure Renilla luciferase activity.

**Bioinformatics and statistical analyses**

Venn diagrams were generated with the online tool available at <https://bioinformatics.psb.ugent.be/webtools/Venn/>. Functional enrichment analyses were performed with DAVID [13,14]. BioRender was used for graphic representations. All statistical analyses were performed with GraphPad Prism. Statistical significance is indicated as followed: *p<0.05, **p<0.01, ***p<0.001.

**SUPPLEMENTARY FIGURES**

***Fig. S1. Impact of lopinavir on HDAg-L secretion and HBsAg level, and cell viability of ritonavir, lopinavir and cobicistat-treated cells.*** **(A)** Dose–response effect of lopinavir (LPV) treatment (48h) on HDAg-L and HBsAg parameters in the Huh7(HiBiT-HDAg-L/S-HBs) cell line as measured by HiBiT detection and CLIA, respectively. **(B)** Cell viability was assessed using the CellTiter-Glo assay following 48 h treatment with ritonavir (RTV), lopinavir (LPV), or cobicistat (CBC) in Huh7(HiBiT-HDAg-L/S-HBs) cells. Mean ± SEM of independent experiments (raw values were first normalized to DMSO control in each experiment). One sample t-test using 100% as reference and Holm-Šidák correction for multiple testing.

***Fig. S2. LDH activity and HBsAg expression in supernatants of PHH co-infected with HBV and HDV, with or without ritonavir treatment.* (A)** LDH release in supernatants was determined to assess cytotoxicity following treatment with ritonavir under the culture conditions described in Fig. 2A. Mean ± SEM of independent experiments (raw values were normalized to DMSO control in each experiment). **(B)** HBsAg was quantified in PHH supernatants by CLIA. Black circles correspond to PHH directly isolated from human liver resections, whereas black squares correspond to PHH purified from the liver of humanized TK-NOG mice (HepaSH®). Mean ± SEM of independent experiments (raw values were first normalized to DMSO control in each experiment).

***Fig. S3. Effect of ritonavir on viral entry in dHepaRG cells.*** Since Ritonavir has been previously reported to inhibit HDV entry in Huh7/NTCP cells with an EC_50_ of 9.1 μM [15], we verified that the presence of ritonavir in PHH supernatants, once diluted to 1/60^th^ corresponding to 0.17 μM of ritonavir, did not interfere with HDV entry into dHepaRG cells. To achieve this, dHepaRG cells were infected with both HBV and HDV in a culture medium containing a dose-response of ritonavir. After 24 h of incubation, cells were washed before adding fresh culture medium. After 6 days of culture, intracellular HDV RNAs were quantified by RT-qPCR and normalized to the PrP housekeeping gene. Results correspond to the mean ± SEM of independent experiments (raw values were first normalized to DMSO control in each experiment). One sample t-test using 100% as reference and Holm-Šidák correction for multiple testing.

***Fig. S4. LDH activity assay in HBV-infected dHepaRG cells treated with ritonavir or lopinavir.*** LDH activity assay was performed to assess cytotoxicity following treatment with ritonavir or lopinavir under the described culture conditions. Results are expressed as percentage of viability relative to negative and positive controls (DMSO and 10% Triton X100, respectively). Mean ± SEM of independent experiments. One sample t-test using 100% as reference and Holm-Šidák correction for multiple testing.

***Fig. S5. Lopinavir alter the biogenesis of HBs in HBV-infected dHepaRG cells.*** Differentiated HepaRG cells were infected with HBV and treated with lopinavir as described in Fig. 2E. Data correspond to the effect of lopinavir on HBV infection. Left panel: intracellular HBV RNAs quantified by RT-qPCR, normalized to human RPL13A expression used as housekeeping gene. Middle panel: HBV genome copies in culture supernatant quantified by qPCR using a standard curve. Right panel: HBsAg level quantified by CLIA in culture supernatant. Mean ± SEM of independent experiments (raw values were first normalized to DMSO control in each experiment). One sample t-test using 100% as reference and Holm-Šidák correction for multiple testing.

***Fig. S6. Effect of lopinavir and ritonavir on S-HBs in Huh7(S-HBs-HiBiT) cells.*** **(A)** Effect of lopinavir treatment (48 h) on S-HBs secretion (HiBiT signal), antigenicity (CLIA; IU/ml), and cell viability (CellTiter-Glo). **(B)** Effect of ritonavir on the same parameters, shown in the same order and format as in panel (A). Mean ± SEM of independent experiments (raw values were first normalized to DMSO control in each experiment). One sample t-test using 100% as reference and Holm-Šidák correction for multiple testing.

***Fig. S7. Ketoconazole alters the antigenicity and the glycosylation profile of S-HBs.*** **(A)** Huh7(S-HBs-HiBiT) cells were treated for 48 h with ketoconazole. S-HBs secretion was quantified via the HiBiT luminescence assay (total secretion), folding was assessed using CLIA targeting the conformational antigenic loop (HBsAg), and cell viability was measured by CellTiter-Glo assay. Mean ± SEM of independent experiments (raw values were first normalized to DMSO control in each experiment). One sample t-test using 100% as reference and Holm-Šidák correction for multiple testing. **(B)** Huh7(S-HBs-HiBiT) cells were treated for 48 h with ketoconazole. Western blot analysis of S-HBs revealed a dose-dependent increase in the proportion of the glycosylated monomeric form upon ketoconazole treatment. The relative abundance of glycosylated versus non-glycosylated forms is shown in the quantification table below the blot.

***Fig. S8. RT-qPCR analysis of ER stress markers in cells treated with ritonavir.*** Expression level of CHOP/DDIT3 (A), and PHGDH, SHMT2, and MTHFD2 (B) was assessed by RT-qPCR following ritonavir treatment. PrP was used as housekeeping gene for normalization. All results are expressed as percentages relative to the DMSO-treated control. Mean ± SEM of independent experiments (raw values were first normalized to DMSO control in each experiment). (A) Friedman test. (B) One sample t-test using 100% as reference.
